# New machine learning method identifies subtle fine-scale genetic stratification in diverse populations

**DOI:** 10.1101/2023.08.07.552391

**Authors:** Xinghu Qin, Peilin Jia

## Abstract

Fine-scale genetic structure impacts genetic risk predictions and furthers the understanding of the demography of populations. Current approaches (e.g., PCA, DAPC, t-SNE, and UMAP) either produce coarse and ambiguous cluster divisions or fail to preserve the correct genetic distance between populations. We proposed a new machine learning algorithm named ALFDA. ALFDA considers both local and global genetic affinity between individuals and also preserves the multimodal structure within populations. ALFDA outperformed the existing approaches in identifying fine-scale genetic structure and in retaining population geogenetic distance, providing a valuable tool for geographic ancestry inference as well as correction for spatial stratification in population health studies.

## Introduction

Fine population genetic structure often arises from social, cultural, physical or geographic barriers and complex demographic history that cannot be easily discriminated. Such structure harbors nuanced genetic diversity that impacts the genetic dissection of complex traits. Up to now, most of the studies mainly focused on genetic ancestry at global scale or on a broad continental level. It remains unclear to what extent the fine-scale genetic ancestry profiles are distinguished and how subtle they are in populations.

Increasing genome-wide sequencing data conducted for diverse populations across the world and the availability of genome-wide data from ancient human samples have revolutionized and advanced the study of fine-scale genetic structure [1–5]. One of the significant progresses is to utilize the geographic information to study the spatial genetic structure [6–10]. Typically, the genetic space significantly aligns with the geographic space or the genetic distance is remarkably correlated with geographic distance (also called isolation-by-distance scenario) [10, 11]. More profoundly, it is reported that the recently originated rare genetic variants tend to be more geographically clustered [3]. These rare variants potentially shape very fine scale population stratifications, leading to false positive results in association studies. On the other hand, culture and socioeconomic differences can shape the genetic structure beyond geography. A typical example is that specific languages and dialects are found to covary with genetic variation on a worldwide scale [12, 13] and in small-scale regions, such as Caucasus [14, 15], the Levant [16], the Amazon [17], and Africa [18, 19]. Recent study found that more subtle dialect-level linguistic differences within a language can influence mating preferences and thus affect genetic population structure [20]. These subtle genetic differentiations have been reported to lead to the health disparities in a series of studies [3, 4, 21]. Despite evidence of population stratification in most human populations due to geographic boundaries and ethnolinguistic differences, we do not yet know how widely the genetic diversity distributes within human populations.

Most commonly used techniques for detecting and correcting genetic structure in association studies are the linear feature representations such as principal component analysis (PCA) [22, 23] and DAPC [24], which use a few top genetic features to represent genetic ancestry. Increasing studies have reported that these linear transformations (PCA and DAPC), however, were not sufficient to fully capture the fine and subtle genomic structure [4, 10, 25, 26]. Further study showed that PCA-based methods of adjustment for population stratification could adequately correct for potential false positives at the large geographical scales, but not so well at the finest scales [3]. Recent studies employed nonlinear dimension embedding, such as t-SNE and UMAP, to visualize population genetic structures [2, 4, 25, 27, 28]. These studies showed that t-SNE and UMAP could reveal cryptic population structure and also preserve the geography of populations [2, 25]. Despite the global positions of clusters are better preserved in some cases, t-SNE and UMAP suffer from overlooked pitfalls. Because t-SNE and UMAP use local notions of distance to construct its high-dimensional graph representation, the size of clusters relative to each other and the distances between clusters are likely to be meaningless [29]. t-SNE and UMAP neglect the local density of data points in the original space, often resulting in misleading visualizations where densely populated subsets of individuals are given more visual space than warranted [30]. Thereby, the reliability of t-SNE and UMAP in inferring spatial genetic structure still needs to be validated both theoretically and empirically.

In this study, we proposed a new machine learning method based on local and global decomposition of genetic affinity which employs nonlinear local discriminant analysis for the dimension reduction. We referred to as Affinity based Local Fisher Discriminant Analysis (ALFDA). ALFDA is based on a weighted graph with edge weights representing the genetic relatedness between two individuals. Like t-SNE and UMAP, ALFDA is a nonlinear method that can capture both local and global structure for multimodal data. But t-SNE and UMAP use stochastic gradient descent to iterate on the graph for embedded space thus may lead to reproducibility problem. In contrast, ALFDA searches for the closest edges on the graph and then preserves the weights between and within populations with singular value decomposition, producing highly stable and reproducible features.

We compared the performance of ALFDA for fine-scale population structure inference with those of PCA, DAPC, t-SNE and UMAP by applying these methods to both simulated and empirical datasets.

## Results

### Performance of methods under simulated spatial scenarios

We designed four spatial scenarios with different spatial structures, including island model, stepping stone model, hierarchical island model and hierarchical stepping stone model (see Methods). We evaluated the performance of all methods for identifying the spatial genetic structures under these four scenarios. Results show that, the most commonly used approach, PCA, is able to impressively retain the global structures of four spatial scenarios. PCA projections intuitively reflect the spatial continuity of genetic variations under the simulated spatial scenarios. Though PCA captures the global structure and clearly distinguishes the aggregates (regional structure) under hierarchical models (hierarchical island model and hierarchical stepping stone model), it is not able to capture the substructures under hierarchical island model and fails to differentiate among populations under the four scenarios (Fig. 1A-D, Fig. S1). DAPC shows similar results with PCA, which is able to capture the global structure but not able to identify the populations within region under all scenarios (Fig. 1E-H, Fig. S1). t-SNE, UMAP, and ALFDA successfully identify the substructure within regions, and are able to clearly discriminate between populations compared to PCA and DAPC (Fig. 1 & Fig. S1). However, among these three outstanding approaches, t-SNE and UMAP seem exhibit global structures that largely mismatch with the first three approaches (PCA, DAPC, and ALFDA) (Fig. 1M-T). PCA, DAPC, and ALFDA distinctly display the variation of populations arranged along a spatial cline under stepping stone model and hierarchical stepping stone model (Fig. 1C, D, G, H, K, L), while t-SNE and UMAP are less capable of retraining the continuity of spatially structured clines (Fig. 1O, P, S, T).

**Fig. 1.**
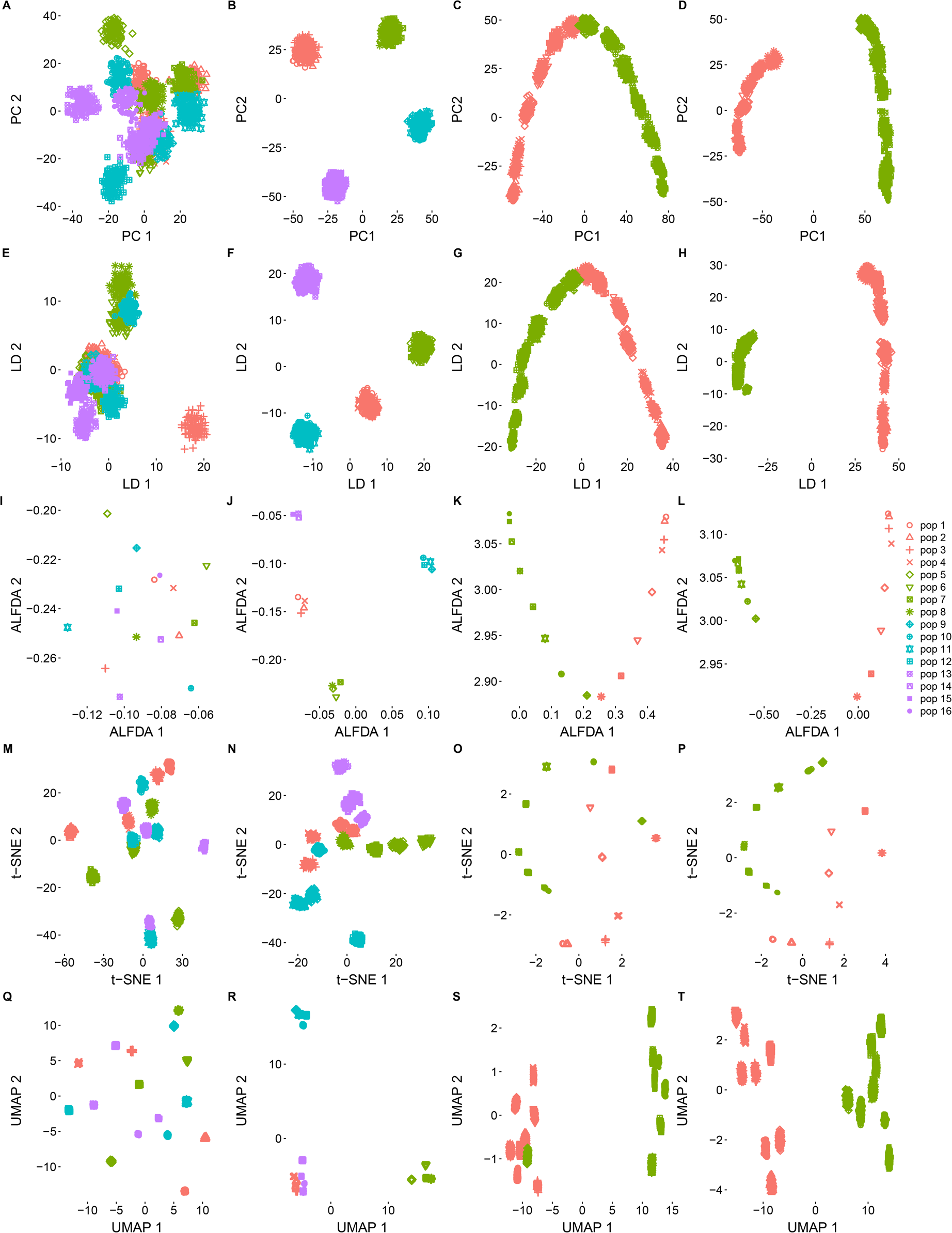
Projection of first two reduced features for four spatial scenarios using five approaches. (A-D), PCA; (E-H), DAPC; (I-L), ALFDA; (M-P), t-SNE; (Q-T), UMAP.

Notably, ALFDA captures both discrete and continuous patterns of genetic variation. It not only shows remarkable ability to identify the local structure, but also keeps the institutive global structure similar with PCA (Fig. 1I-L). Further investigation into the correlations between the reduced features shows that, among the five approaches, only PCA and ALFDA always keep high significant correlations between their first two reduced features in all three scenarios (hierarchical island model, stepping stone model, and hierarchical stepping stone model) (Fig. S2).

ALFDA, t-SNE and UMAP exhibit higher power than PCA and DAPC in differentiating populations under four spatial scenarios (Fig. S3). It is worth mentioning that, ALFDA not only has the highest overall accuracy among the five approaches in discriminating among populations (Fig. S3), but also has the highest correlation between the *F_ST_* and the Euclidean distance of the genetic features under the isolation-by-distance model (Fig. S4). Compared with the existing approaches, and combining the visualization results above, ALFDA substantially increases its power to detect subtle levels of genetic differentiation, outperforming PCA and DAPC in population discrimination, as well as t-SNE and UMAP in spatial pattern preservation.

### Population structure of 1000 Genomes

Application in the global human populations (1000 Genomes dataset) highlights the advantages of ALFDA in identifying latent population structures that were not found using the traditional approaches. Representations of population structure from PCA, DAPC, and ALFDA projections intuitively keep the same well known “V” shaped global population structure (Fig. 2A-C), while population structures represented by t-SNE and UMAP do not retain such a triangle global structure. Instead, t-SNE presents continuous aggregates with several subpopulations visible within aggregates and UMAP presents discrete aggregates with some overlaps between European, African, Asian and American populations, respectively (Fig. 2D-E).

**Fig. 2.**
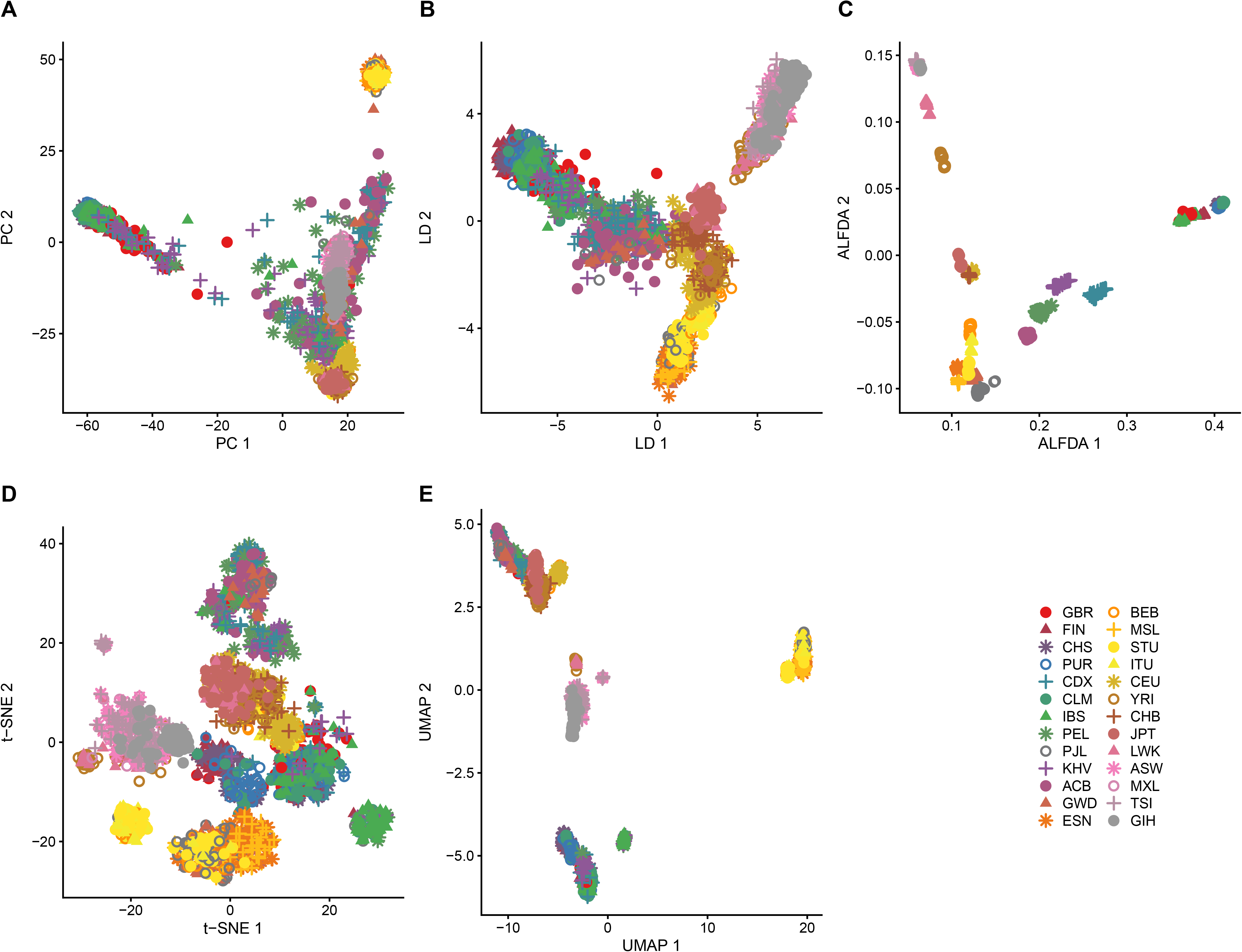
Population structure of 26 populations from 1000 genomes data. Representation of population structure using PCA (A), DAPC (B), ALFDA (C), t-SNE (D), and UMAP (D).

Generally, PCA and DAPC present the similar continuous patterns of genetic variation in a triangle with vertices. PCA and DAPC cluster the individuals into three aggregates, among which individuals from Asia, Europe, and Africa can be roughly discriminated (Fig. 2A). However, both PCA and DAPC fail to identify populations between countries (Fig. 2A-B).

In contrast, ALFDA exhibits considerable continuous patterns of distinct aggregates (Fig. 2C). It is worth noting that ALFDA clearly clusters all the individuals into 25 populations at the two-dimensional plane, and reveals more detailed subtle substructure within countries (e.g., LWK, YRI, JPT, BEB, ITY, STU, GWD, PJL, PEL, KHV, CDX, IBS, GBR; Fig. 2C).

Furthermore, ALFDA identifies three distinct sub-clusters within Punjabi in Lahore, Pakistan (PJL) and Chinese Dai in Xishuangbanna (CDX), and a continuous cline in Kinh in Ho Chi Minh City, Vietnam (KHV) (Fig. 2C). In addition, ALFDA shows close genetic affinities between Toscani in Italy (TSI), Mexican Ancestry in Los Angeles (MXL), Gujarati Indian in Houston (GIH), and African Ancestry in Southwest US (ASW) as well as close genetic affinities between Han Chinese in Beijing (CHS), Puerto Rican in Puerto Rico (PUR) and Colombian in Medellin, Colombia (CLM). These results, which may reflect specific demographic histories in these populations, are hardly drawn from other approaches (Fig. 2).

Although t-SNE presents 9 impressive clusters, with each cluster also exhibiting several overlapping substructures (Fig. 2D), the clusters and sub-clusters from t-SNE present complex distribution that are distinct from other approaches. For example, African Caribbean in Barbados (ACB), Peruvian in Lima (PEL) and CDX, who are from Africa, America, and east Asia, respectively, are clustered together (the top cluster at Fig. 2D). However, this cluster is a unique cluster that other approaches do not present (Fig. 2). Meanwhile, the identities of some aggregates in t-SNE projection are in line with ALFDA result (Fig. 2C-D). For example, Mexican Ancestry in Los Angeles (MXL), Toscani in Italy (TSI), African Ancestry in Southwest US (ASW), Gujarati Indian in Houston (GIH), Luhya in Webuye (LWK), and Yoruba in Ibadan (YRI), which are also identified to have close affinities by ALFDA, are clustered together with high overlap in t-SNE projection (Fig. 2D).

UMAP clearly distinguishes 4 large aggregates, with each presenting several sub-aggregates that have the same identities with ALFDA and t-SNE projections (Fig. 2E). In line with the results of ALFDA and t-SNE, MXL, TSI, ASW, GIH aggregate together as well (Fig. 2C-E). However, UMAP presents global discrete aggregates with some aggregates clustering some distant populations therefore are difficult to interpret (Fig. 2E). For example, three populations, Punjabi in Lahore, Pakistan (PJL), Chinese Dai in Xishuangbanna (CDX), and Kinh in Ho Chi Minh City, Vietnam (KHV), which present subtle substructures in ALFDA, are all divided into two separate clusters in UMAP projection (Fig. 2D). Furthermore, UMAP projection keeps a similar global triangle structure with PCA, DAPC and ALFDA. However, the identities of aggregates and sub-aggregates in UMAP do not match the similar continuous distribution pattern compared to PCA, DAPC and ALFDA (Fig. 2A-C, E). Among all these methods, ALFDA performed the best in differentiating 26 populations (Fig. S5).

Overall, the identifiability of these five approaches differs largely. Nevertheless, ALFDA shows the most outstanding performance in presenting the discriminable continuous patterns of genetic variation both locally and globally.

### Small scale and fine scale population structure of the world largest population

We identify the individuals of the world largest population, Chinese, at the province level (small scale within nation, including 19 provinces, 4 municipalities, and 1 autonomous region) using CONVERGE dataset (China Oxford and Virginia Commonwealth University Experimental Research on Genetic Epidemiology) [54]. PCA result indicates there is no discernable geographical structure for populations whether from south to north or from west to east (Fig. 3 & S6). However, a weakly recognizable South-North structure is found at province level in DAPC, though populations among provinces are still not distinguishable (Fig. 3 & S6A-B). Compared to PCA, DAPC makes an improvement in identification of South-North populations (Fig. 3A-B & S6C). Although there is a South-North geographical structure, most of the populations in DAPC projections are still not differentiable (Fig. 3A & S6C).

**Fig. 3.**
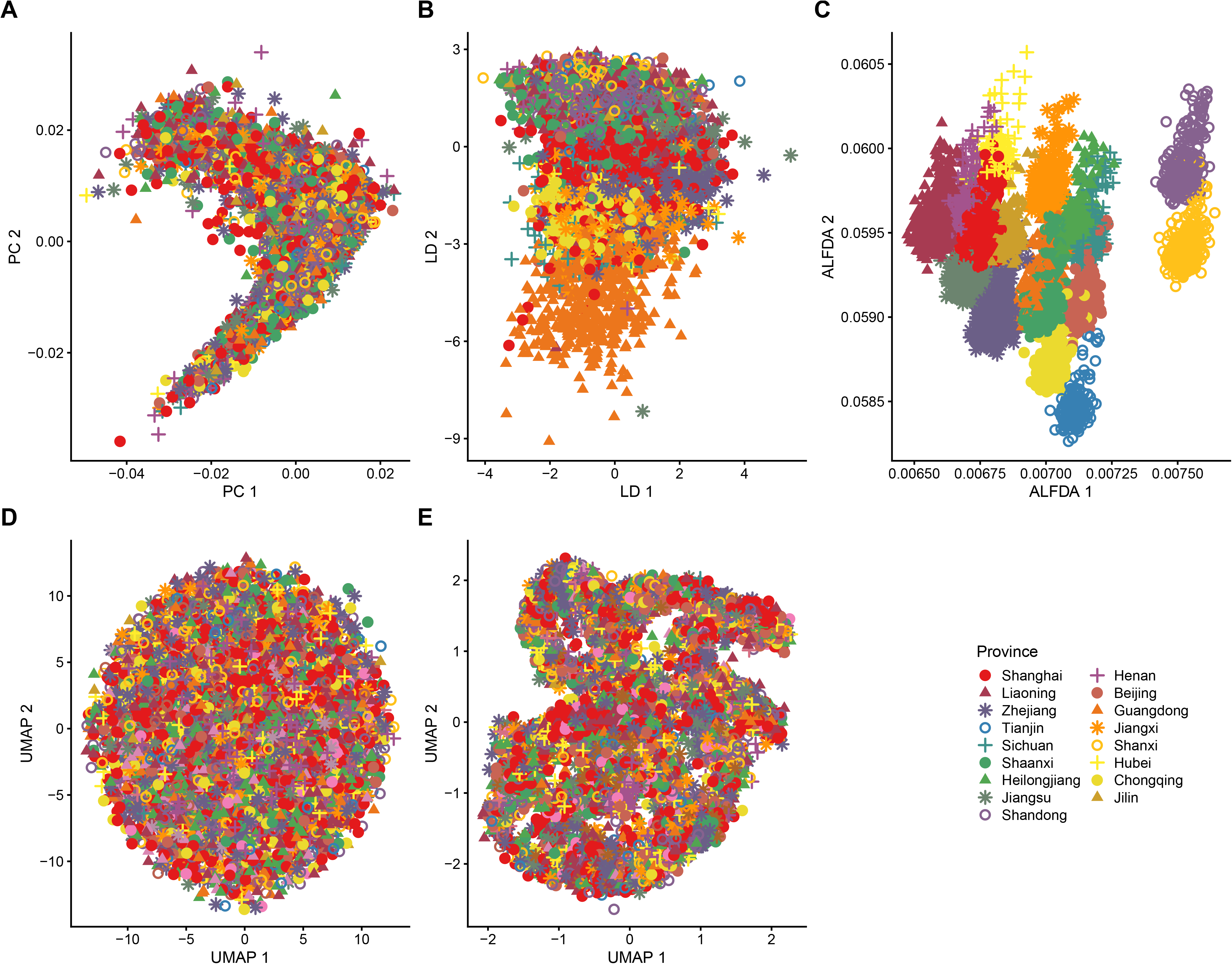
Population structure of Chinese individuals from 17 provinces represented by (A) PCA, (B) DAPC, (C) ALFDA, (D) t-SNE, and (E) UMAP. Individual labels were assigned based on their self-reported birth locations (based on the ID).

In contrast, ALFDA representation of populations presents more clear and subtle structures (Fig. 3C & S6D). Compared to PCA and DAPC representation, ALFDA representation provides more precise population structure identifiability (Fig. 3A-C).

More clear subtle genetic structures are revealed after the genetically distant populations (i.e., Hunan, Hebei, Fujian, Gansu, Anhui, and Hainan, Fig. 3) are hidden. Compared to other approaches, ALFDA identifies deep genetic relatedness and complex demographic histories between populations as indicated from the spatial patterns. We observe a close relatedness between adjacent provinces, e.g., there are a small amount of mixtures between Shandong province and Shanxi province (Fig. 3 & S6-7). We also observe that geographically distant provinces but high-developed provinces, such as Beijing, Chongqing, and Tianjing, aggregate closely (Fig. 3 & S6-7). Moreover, investigation of the genetic distribution at the fine scale shows that there is a large amount of mixture among highly developed provinces and underdeveloped provinces. One of the most developed cities in China (provincial administrative city), Shanghai, is the most mixed populations among all the provinces (Fig. 3 & S6-7; Appendix 2). Shanghai not only has genetic mixtures from the adjacent provinces such as Jiangsu and Zhejing, but also has a large amount of mixtures from geographically distant provinces such as Henan, Jilin and Liaoning (Fig. 3 & S6-7).

t-SNE clearly distinguishes 8 populations at the province level except that, the rest of populations are mixed together without a clear differentiation (Fig. 3E & S6E). However, t-SNE fails to identify clear substructure compared to the above three approaches (Fig. 3E & S6E).

UMAP only successfully discriminates Guangdong among other provinces at the province level (Fig. S6F). When looking into the large aggregations, aggregations present several continuous substructures. However, these substructures are chaotic without any identifiability (Fig. 3F & S6F).

Among all these five methods, ALFDA has the highest discriminatory power to identifying populations from different provinces, significantly outperforming the other four methods (Fig. S8). In summary, compared to PCA, DAPC, t-SNE and UMAP, ALFDA can further identify subtle subpopulations within the country, despite their geographic adjacency.

### Subtle genetic structure reconstructs the geographic origin of ancient British individuals

By apply all five methods to an ancient British population, four approaches (PCA, DAPC, t-SNE, and UMAP) presents the similar genetic structure pattern where all populations are clustered together and it is impossible to distinguish the Channel Islands and the Wales populations from the other four populations. On the contrary, ALFDA clearly discriminates these two populations among other four populations which present some sub-structures (Fig. S9). In addition, PCA could discriminate some individuals from Netherlands, France and Scotland (Fig. 4B & S9A). These results indicate that there is high genetic admixture among individuals from England, Netherlands, Scotland and France. When looking into the genetic structures of these four populations, PCA, DAPC, t-SNE, and UMAP still cannot distinguish these four populations clearly (Fig. 4). Most of the individuals are clustered together centred by England. Instead, ALFDA successfully differentiates four populations and presents clear fine-scale genetic structure (Fig. 4D). Spatial genetic structure of four population projected by ALFDA is highly concordant with the geographic origin of these populations (Fig. 4D, PC1 vs. latitude r=0.967, p-value=2.67e-14). The first reduced feature corresponds to the north to south UK geographic axis, which separates the populations into four distinct clusters representing their geographic locations. These results indicate that ALFDA not only improves the performance of discriminatory power but also improves the performance of reconstructing the geographic origin of individuals.

**Fig. 4.**
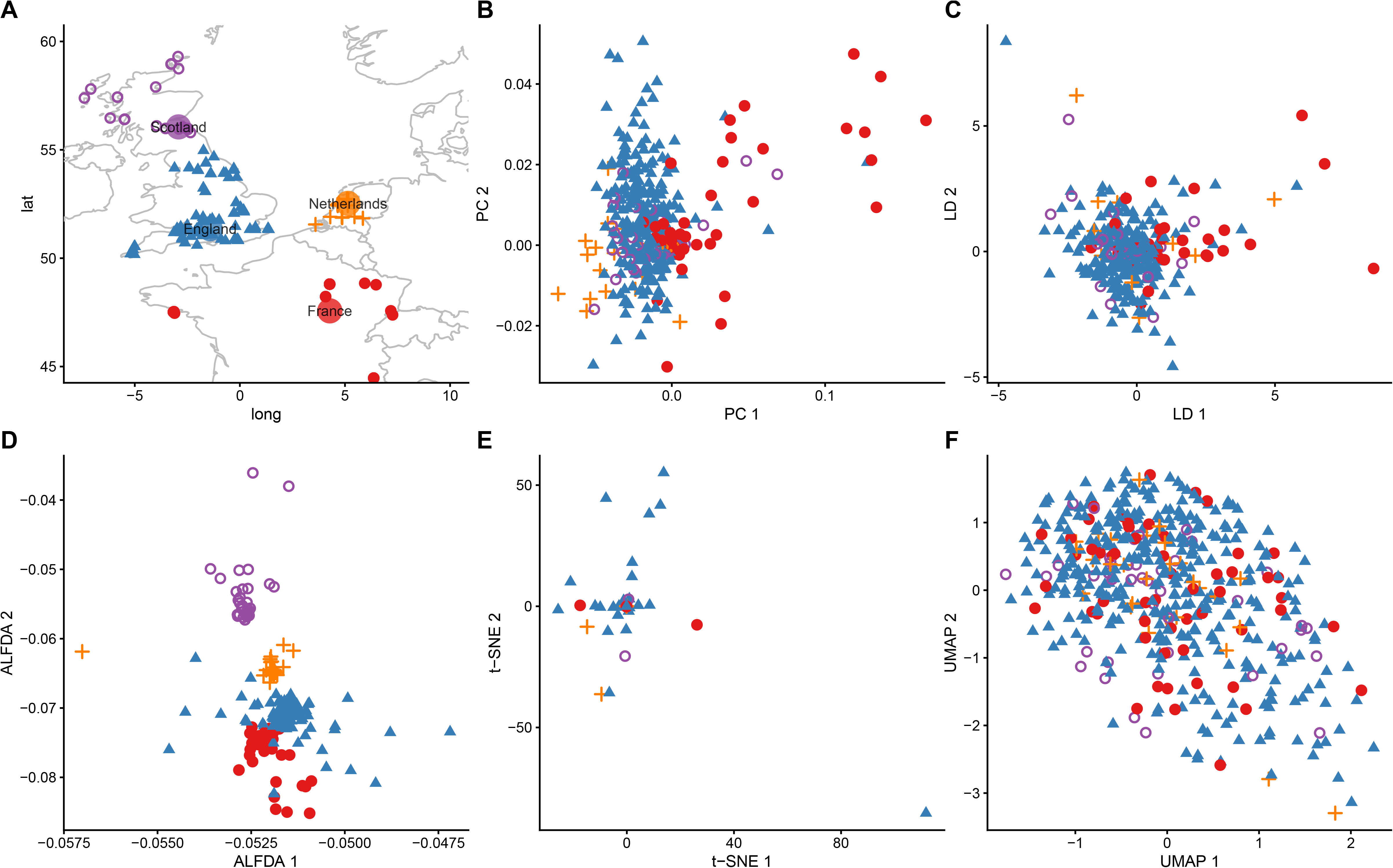
Population structure of four populations from Britain and Europe. (A) The geographic distribution of individuals 4000 BCE-43 CE. Population genetic structure projected by PCA (B), DAPC (C), ALFDA (D), t-SNE (E), UMAP (F).

## Discussion

Heretofore, human population genetic structure is fairly well understood at the global scale or on a broad continental level, but the fine-scale population genetic stratification within some unrepresented local populations remains poorly understood. In this study, we proposed a new machine learning approach, ALFDA, for fine-scale genetic structure inference. We tested the performance of ALFDA for fine-scale population structure inference using both simulated genetic data and empirical datasets. Simulation studies showed that ALFDA significantly improved the accuracy of identifying genetic structure compared with PCA and DAPC, especially in identifying the substructures within regions. In addition, ALFDA did better in recapitulating the spatial position of individuals compared to t-SNE and UMAP, where ALFDA showed a stronger correlation between the *F_ST_* and the Euclidean distance of the genetic features under the isolation-by-distance model. Empirical population data analysis showed that ALFDA performed better than PCA, DAPC, t-SNE and UMAP in identifying fine-scale population genetic structure in diverse populations. When applied to the 1000 Genomes data, all five approaches presented a similar “V” structure. But ALFDA identified new genetic structures within populations that were not detected by other approaches. When applying five methods to Chinese population dataset, PCA cannot distinguish the genetic gradient across populations. DAPC improved discrimination of North-to-South genetic structure compared to PCA, while t-SNE and UMAP presented highly chaotic structure that were not discernable. In contrast, ALFDA not only presented the geographic clines of genetic variation, but also presented the discernable boundary between populations. Analyses of ancient British individuals indicated that ALFDA significantly improved the identification of fine-scale genetic structure and correctly reconstructed the geographic origin of individuals that would not be apparent via the other methods. Taken together, this study indicates ALFDA, which has impressive power to identify fine-scale genetic structure while preserve the multimodal structure of the individuals, can serve as a useful alternative tool over PCA, DAPC, t-SNE and UMAP for many population genetics studies.

One key aspect of genome-informed precision medicine is the accurate detection of ancestry to understand its impact on disease susceptibility and the outcomes of therapies. Conventionally, population geneticists often relied on model-based methods (STRUCTURE, ADMIXTURE) or linear dimension reduction methods to study population genetics. However, model-based methods generally assume that the allele frequency in a population is constant. This sometimes violates the realistic scenarios. On the other hand, PCA projects genomic data onto a low-dimensional representation that captures as much variance as possible. The distances between populations in a PCA plot can be interpreted in terms of times to the most recent common ancestors [57]. However, PCA usually ignores the within population differentiation and tends to obscure finer-scale patterns of population structure [10, 58]. Although DAPC considers the within genetic structure, it weights the between population affinity using the centroid/mean of the population features, inevitably distorting the multimodal genetic structure. Recently, studies tried to visualize the genetic structure of ancient DNA samples combined with modern and contemporary populations using t-SNE and UMAP [59]. Nevertheless, the distance between points produced by t-SNE and UMAP does not represent the real distance between two samples [29]. t-SNE and UMAP representations can also contain arbitrarily small distances between points. The positions of connected clusters in t-SNE and UMAP may be flipped or rotated relative to each other when carrying out multiple runs with identical parameters. This limitation leads to misleading visualizations where the apparent size of a cluster largely reflects the number of points in the cluster rather than its underlying heterogeneity [30].

In our study, we show that PCA and DAPC generally produced similar global genetic structure but they still cannot identify substructure within continent (region) whether in simulation studies or empirical data analysis. Although t-SNE and UMAP have high discriminatory power in simulation studies, they performed poorly in real data analysis. Furthermore, the UMAP and t-SNE does not by itself decide when it has converged but simply does ’maxSteps’ iterations. In contrast, ALFDA outperformed the other methods both in simulation studies and real data analyses in terms of the discriminatory power and geogenetic relation. ALFDA identified subtle genetic structures in diverse populations that were not discovered by the other methods.

PCA has been a workhorse for describing population structure for decades. Previous study reported that PCA only partially identifies population clusters and does not separate most populations within a given continent, such as Japanese and Han Chinese in East Asia, or Mende and Yoruba in Africa [60]. t-SNE and UMAP, on most of the genomic data analysis cases, cannot identify more fine-grained population clusters but produce chaotic clusters or aggregations such as presented in [4, 25, 27, 28]. In our study, ALFDA not only presented the geographic clines of genetic variation, but also presented the discernable boundary if populations are not recently mixed. In empirical datasets, sampled individuals were collected from geographically or culturally distinct groups, ALFDA arguably provided a more detailed representation of the relationships among study participants compared to other methods.

Rare alleles likely originate from recent mutations and are geographically clustered around the location where the mutation initially arose. Thus, they can be highly informative about gene flow and the fine-scale population structure [61]. Typically, when analysing the genetic structure, one common step to eliminate errors is to remove the rare variants with a MAF below some specified threshold (e.g., 0.01). Recent study revealed that both model-based and multivariate analyses infer less distinct clusters when using larger MAF which is supposed to have removed alleles originated more recently [62, 63]. This is an important concern when analysing ancient DNA datasets. Due to the uncertain genotyping quality in ancient genomes, we chose a MAF of 0.01 for analysis, which indicates alleles only exist in appropriately 5 individuals in this study. These low frequency alleles would include important information about the mutation and gene flow, thus informative about fine-scale population genetic structure. However, PCA, DAPC, t-SNE and UMAP all failed to reveal clear genetic structure of five populations when analysing ancient genomes (Fig. 4). Instead, ALFDA not only uncovers fine-scale genetic structure, but also correctly recapitulates the individual geographic origin.

The advantages of ALFDA originate from the strategies for characterizing the genetic affinity of individuals with respect to the genotypes they have, both within and across populations. Genome wide affinity has been already used with respect to DNA alignment [64, 65], phylogeny analysis using genome distance (delta distance) [66] and genome clustering [67]. Our method introduces a new similarity measure to compute the genetic distance and then uses the KNN graph to estimate the genetic affinity (weight) between individuals. This adequately reflects the contribution of untyped markers to relatedness. It thus provides improved ancestry maps by accounting for a higher percentage of explained variance in ancestry.

Even though rapid progress has been made on the methodological development, many fundamental challenges exist for fine-scale population structure identification. Many of these challenges originate from the difficulty of recapitulating the complexity of human population structure with mathematical models. On the one hand, many models predefine the discrete populations as input or assume temporally continuous individual as the input for analysis. The errors of these analyses can be induced when the discrete populations violate the model assumption. On the other hand, non-parametric models don’t need to predefine any assumptions. Many of these types of methods focus on extracting the observed population structure in a representation that is useful for interpretation, but without explicitly modelling the historical processes that shaped these patterns. We also believe the mathematical explanations and spatial genetic modelling process by ALFDA in our simulation studies provide informative interpretation on how we can recapitulate the spatial pattern of genetic structure under different scenarios. However, explicitly studies that consider wide fine-scale genetic scenarios are still needed to model the historical processes that shaped fine-scale genetic structures.

In summary, this study proposed a new machine learning method, ALFDA, and proved its remarkable performance in identifying fine-scale population genetic structure. ALFDA could preserve the multimodal genetic structure within diverse populations, overcoming the impact of minor allele frequency defects in ancient DNA studies, and correctly distinguishing the fine stratification of genetic admixtures. This study reinforces our understanding of how nuanced the fine-scale population structure is and provides a useful tool toward fine-scale monitoring of population health.

Without any doubts, the study of fine-scale structure has been an exciting frontier of contemporary population genetics, with extensive progress and continued promise. As this work continues, we will begin to more fully understand the processes that shape fine-scale structure in humans, and have a fuller perspective on human origins. Of broader relevance, this progress also provides guidance for studying other species with highly dynamic population histories, and the methods we introduced in this study are useful for applications outside of humans.

## Methods

Our affinity-based local and global structure preservation algorithm is constructed following three steps (Fig. 5). First, we construct the Shannon similarity matrix using the genetic relatedness among individuals. Second, we estimate the genetic affinity (weights) based on the Shannon similarity matrix from step 1 using *k*-nearest neighbours (KNN). Third, we conduct dimension reduction on the affinity matrix to preserve the local structure while embedding the multimodal spaces using nonlinear local Fisher discriminant analysis. The nonlinear local Fisher discriminant analysis makes use of the nonlinear mapping to produce the localized global pairwise affinity that maximizes between-class affinity and minimizes within-class affinity while preserves the multimodal structure. Therefore, our method is also called affinity based non-linear local Fisher discriminant analysis (hereafter referred to as ALFDA).

**Fig. 5.**
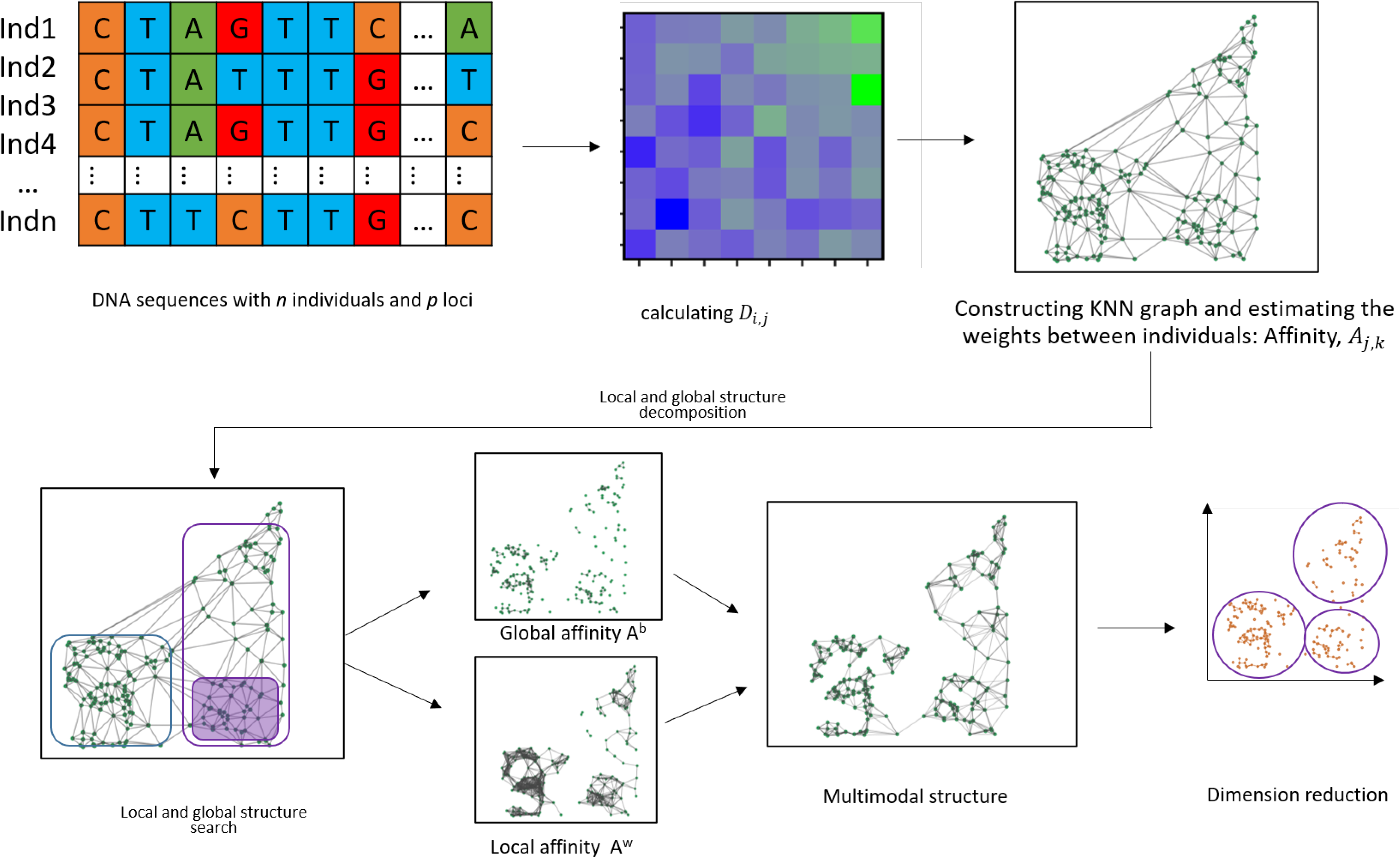
The schematic workflow of ALFDA algorithm. The input genotype matrix (A) with individuals in rows and markers in columns imputed from vcf file is used to calculate the genetic similarity (distance) matrix representing the genetic relatedness among individuals (B). (C) Similarity network represented as a graph is constructed using KNN graph, and the edges (weights) are represented as genetic affinity between individuals. The local and global structure are identified (D) and then the local and global affinity matrix are computed (E), thus the multimodal structures are preserved (F). The final reduced features are produced for genetic structure inference through the dimension reduction. D-G is implemented following KLFDA algorithm.

Below we explain the details of the main steps in use to infer population structure.

### Constructing individual genome-wide similarity

Genetic association studies, such as disease related gene identification, ancestor inference and forensics, are much dependent on genetic affinity or relatedness that reflects the population stratification. In this study, we propose to use Shannon similarity to construct individual genetic distance, which estimates the genetic similarity between individuals and considers the weight of the derived alleles/reference alleles in the individual amidst all the studied samples. Our approach is analogous to an identity-by-descent (IBD)-based genetic relationship matrix which incorporates mutual shared information to infer finer-scale genetic relationships underlying the structure or demographic history of the study population. Unlike most other relatedness measures that ignore the weight of genetic variants in the samples, our approach computes the genome-wide genetic relatedness between individuals considering the weight of the derived alleles/reference alleles in each individual, explicitly modelling the shared genealogical relation that related to every other individual in a population.

Taking the genome wide variants as an example, we demonstrate how to derive the genetic similarity from single-nucleotide polymorphism data.

To compute genomic distance between two individuals, we defined a genomic dissimilarity matrix across all pairs of SNPs based on Shannon entropy. This genomic dissimilarity corresponds to the distance between all the genome wide genetic variants of two individuals, which considers multi-locus-based independent genotype information across the whole genomes of individuals, in other words, the essential information of combinations of genotypes.

We have *N* samples with *L* loci as an *N*×*L* matrix *X*. The genotype of the individual *i* and *j* coded in terms of the number of copies of the reference allele at locus *l* is *x_i,l_* and *x_j,l_*, respectively. The total number of copies of the reference allele in all individuals is 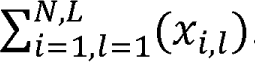.

The Shannon information of individual *i* is,

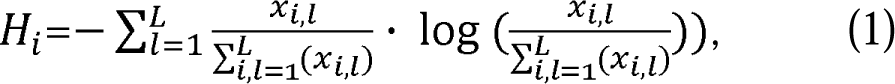

and the Shannon information of individual *j* is,

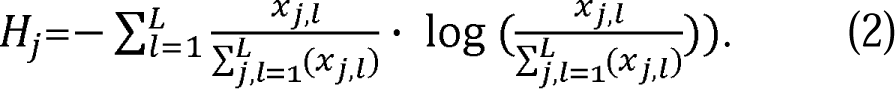

If there is no hierarchical structure within populations, the allele frequency of locus *l* at the pooled individuals is,

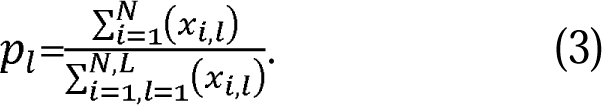

The Shannon information of the pooled individuals is,

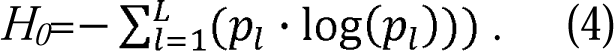

The genomic dissimilarity between individual *i* and *j* is,

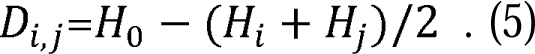

If there are at least two levels of spatially hierarchical structures (e.g., populations, regions), the weight of entity that comprises of two individuals *i* and *j* among all individuals is,

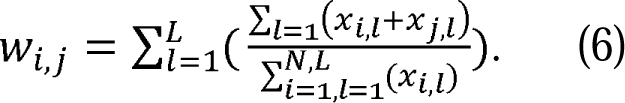

The Shannon information of the entity that comprises of two individuals *i* and *j* is,

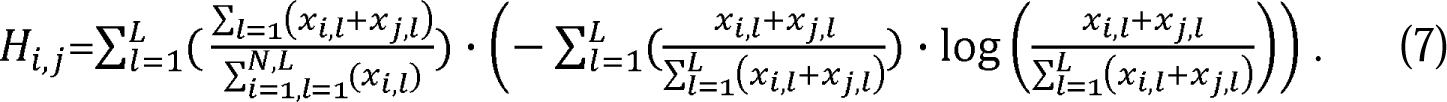

The genomic dissimilarity between individual *i* and *j* is,

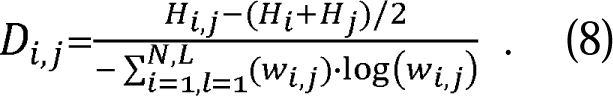

The genomic similarity between two individuals is,

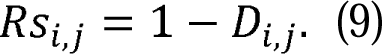

Two individuals that have the same DNA sequence will have a similarity score of 1. The datasets used to calculate similarity can be extended to other markers and data types, such as DNA methylation, mRNA expression, and proteomic profiles.

### Estimating the local genetic affinity

We constructed the genetic affinity between individuals based on the concept of the *k-nearest neighbours graph* (*k-NNG*) [31–33]. The key idea of constructing affinity matrix is to assume the individuals are connected by a similarity network G = (V, E). The nodes V represent the individuals (*x*_1_, *x*_2_, …, *x_n_*), and the edges (*E*) represent the genetic similarity between individuals. The *K-NNG* network captures the local genetic affinity from the underlying genetic features by searching the closest individuals and then constructing the weight between them.

Let *Rs* (*x_i_*, *x_j_*), be the genetic similarity between sample *i*, *j* calculated from equation (9). **W** is an *n* × *n* similarity matrix representing the edge weights. The weight (W*_i,j_*) of the similarity between sample *x_i_*, and *x_j_* is determined by a scaled exponential similarity kernel,

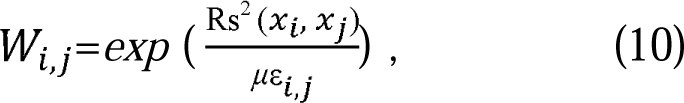

where μ is a hyperparameter that can be empirically set. ε_i,j_ is used to scale the data based on the data type. ε_i,j_ can be estimated as follows,

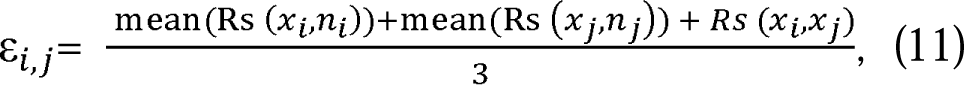

where *n_i_* represents a set of *x_i_*’s neighbours including *x_j_*in subspace. The number of neighbours were searched based on the neighbourhood size *K*. mean Rs(*x_i_,n_i_*) is the average value of the similarity between *x_i_* and each of its neighbours.

The local affinity matrix *A^us^_g,ij_*, is determined by the weight of the similarity *W_i,j_* calculated from equation (10), through the *k*-nearest neighbour (*KNN*) [31–33]. Let *n_i_* represent a set of *x_i_*’s neighbours including *x_i_* in G. Given a graph, G, the local affinity *A^us^_g,ij_* can be calculated as,

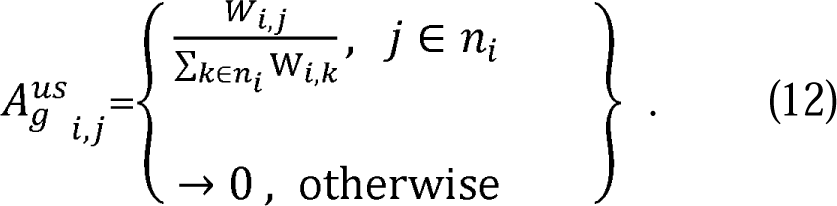

*A^us^_g,ij_* emphases local affinity to the *K* most similar samples. *K* is a defined neighbourhood size used to construct the neighbour search.

There are two parameters, local variance μ, and the number of nearest neighbour *K* described above. μ is a hyperparameter for the scaled exponential similarity kernel used to conduct the actual affinity calculation. *K* is an important parameter to construct the local affinity. In practice, the optimal *u* value can be estimated by the variance of *n*-dimensional structure from training samples, which we recommend in the range of [0.5-1]. Optimal *K* can be found through grid search.

### Non-linear Local Fisher Discriminant Analysis for dimensionality reduction

Although we have obtained the genetic affinity matrix using *k*-nearest neighbour above, the genetic affinity can overlap different structured populations because the genetic affinity, ^us^ is still a *n*l7*n* matrix. The above affinity is learned through unsupervised process. Here, we carry out the supervised learning through the non-linear local Fisher discriminant analysis of dimension reduction to optimally preserve the neighbourhood structure of the pairwise affinity matrix [34], and to get the local within-class scatter matrix and between-class scatter matrix.

Kernel Local Fisher discriminant analysis (KLFDA) is a kernelized version of local linear discriminant analysis (LFDA) used to carry out dimension reduction. PCA does well in capturing the global variances of the data, but not so well when analyzing the local variances within populations. LFDA is good at capturing both the local variances and between population variances. Using the kernel transformation technique, non-linear mapping is learned in KLFDA and the power of discriminating nonlinear separable samples is significantly improved [35]. The diagonal values of the kernel distance will be distorted if the number of features is significantly larger than the number of samples (p >> n) [36]. Thus, kernel distance cannot fully model the genetic relationships between individuals from datasets containing millions of features. Contrast to kernel distance, the genetic affinity fully preserves the non-linear information between individuals, possessing many advantages than kernel matrix. Therefore, instead of using the kernel local fisher discriminant analysis, we adopt the genetic affinity matrix in non-linear local fisher discriminant analysis. The steps to construct the non-linear local Fisher discriminant analysis are described in supplementary materials.

### Posterior probabilities of individual membership

A typical probabilistic model based on Bayes’ theorem is usually used to assign and predict new individuals to the potential groups. The PCA, t-SNE and UMAP do not provide the probabilistic model for membership assignment. We can make use of the reduced features to approximate the likelihood of each individual belonging to the potential labelled populations. Here, the probability of an individual or ancestry can be estimated given the prior probability of the known individuals *X_c_*.

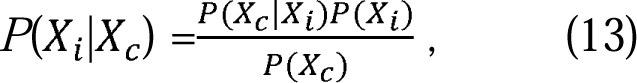

where *P*(*X_i_*|*X_c_*) is the probability of the reduced features for the *i*-th individual, _i_, assigned to any group *c* given that individual’s genetic feature, *X*_c_, having their memberships in *c* group. *P*(*X_i_*|*X_c_*) is the likelihood of *X_c_* occurring in *c* group given *X_i_* is true. *P*(*X*_c_) and *P*(*X*_i_) represent the likelihood of the occurrence of *X_c_* and *X_i_* independently.

We use the reduced features from ALFDA to estimate the probabilities of individuals assigning to the potential known populations based on Eq. (10). The performance of the reduced features for each approach (PCA, DAPC, ALFDA, t-SNE, UMAP) was also evaluated in the following simulation studies following [37].

### Simulation studies

To test the ability of our method to infer the correct population structures and to compare the performance of our method to the existing traditional machine learning approaches (PCA, DAPC) and the latest state-of-art machine learning approaches (t-SNE, UMAP), we simulated four spatial scenarios that differ in hierarchical structure.

We simulated four scenarios using the coalescent-based genetic data simulator fastsimcoal26 [38], which are the island model, the stepping stone model, the hierarchical island model, and the hierarchical stepping stone model. We simulated 16 populations for each model and each population consisted of 1000 diploid individuals. The island model (including hierarchical island model) and stepping stone model (including hierarchical stepping stone model) differ in the composition of aggregates (regions) and migration mode. The island model and hierarchical island consist of four regions with each region comprising 4 populations. The stepping stone model and hierarchical stepping stone model consist of two regions with each region comprising 8 populations. We simulated 44 independent chromosomes (22 pairs) with 100 Kb DNA sequences per chromosome assuming a finite mutation model with a constant mutation rate of *u*=1×10^-8^ per base pair per generation and a recombination of *r*=1×10^-8^ per base pair per generation for all scenarios which are typical of humans. In the case of the non-hierarchical scenarios (island model and stepping stone model), we assumed the migration rates between populations to be 0.001. In the case of the hierarchical models (hierarchical island model and hierarchical stepping stone model), migration rates between pairs of populations within regions were 0.001 and migration between populations from different regions were 1×10^-4^. The four spatial scenarios and their parameters used in the simulations are presented in Table S1.

We sampled 100 diploid individuals from each population for each scenario. Each scenario generated more than 27, 000 polymorphic sites. We did a Linkage Disequilibrium (LD) based SNP pruning with the threshold of r^2^ = 0.2. We randomly selected 10, 000 sites (biallelic) per scenarios for downstream analysis.

### Performance of five approaches in identifying population structure

We converted the arlequin files simulated from fastsimcoal26 into *vcf* format using PGDSpider 2.1[39]. Then the genotypes of each scenario were imputed using SNPRelate package [40]. To compare the performance of our methods in population structure inference with the traditional methods, we conducted principal component analysis (PCA) and Discriminant Analysis of Principal Components (DAPC) as the benchmarks. PCA was implemented using SNPRelate [40]. DAPC was implemented using the first 20 PCs from the PCA results via *lda* function from *MASS* package [41], which is initially employed by *dapc* function in *adegenet* package [42]. The top reduced features from PCA and DAPC were used for population structure visualization. In addition, we calculated the pairwise *Fst* [43] between populations under each scenario as the reference (Appendix 1).

We carried out ALFDA to the genotypes of individuals with different local variance (*u*), the regularization parameter θ, and the number of nearest neighbors *K*. We tuned these parameters to determine the best model through a grid search for each scenario. The recommended default θ value is 0.001. We tuned the parameter *u* from 0.05 to 10 (0.05, 0.25, 0.5, 0.1, 1, 5, 10). Through parameter tuning, we found the optimal *u* values for four models are between (0.5, 5).

We employed the Barnes-Hut implementation of t-SNE [44] and UMAP [45] on four spatial scenarios. In case of t-SNE, there are two important parameters determining its performance, *perplexity*, which is used to balance the diversity between local and global structure, and maximum iterations (*iter*), which is the maximum number of iterations for the optimization [46]. We tuned perplexity from 5-50 (5, 10, 15, 30, 50) and the maximum iterations from 100 to 2000 (100, 500, 1000, 2000) based on the suggestions from the original paper [46]. The optimal parameter combination was determined by the quality of projection and KL divergence value [46].

In case of UMAP, the key parameters affecting the robustness of the model are the number of nearest neighbours (controls how UMAP balances local versus global structure in the data) and minimum distance (the minimum distance that points are allowed to be clumped in the low dimensional representation) [47]. We tuned the number of nearest neighbours from 20 to 99 (20, 40, 60, 80, 99) and the minimum distance from 0.05 to 0.99 (0.05, 0.1, 0.5, 0.99) based on the previous reported cases [2, 25, 37, 47]. The criterion to determine the best parameters for UMAP projection seems uncertain, depending on the purpose of identifying substructure of population or the separability of population. We identified the best-performing parameters via the largest discrimination success-number of successful discrimination cases (or assignment accuracy associated with the population labels) following [37].

We aimed to compare two important properties of our approach with the current state-of-art machine learning techniques: the ability to identify population structure, and the power to retrain the geography or origin of the individuals. To do so, three steps were conducted to show their performance. First, the top (first three) features from above analyses were obtained for population structure visualization under the optimal parameters. Scatter plots of the reduced features for each scenario from each approach were used to compare the quality and ability as the representation of population structure under the corresponding scenario. Second, the accuracies of these five approaches in discriminating and predicting the populations and regions were tested used their first 3 reduced features via 10-fold cross validation through the unbiased random forest classifier [48]. Cross validation was repeated ten times and the overall accuracy was used as the indicator of performance [49]. Third, we estimated the correlation between population genetic fixation (*F_ST_*, [43]) and the Euclidean distance >0 between cluster centroids under isolation-by-distance scenario (stepping-stone model), in which genetic differentiations between populations should increase with geographic distance. We calculated the Euclidean distance between each pair of population centroids based on the first three reduced features for each method. We then ran the Mantel test [50] with 9,999 permutations to estimate the correlations between the *F_ST_* and the Euclidean distance of the genetic features under stepping-stone model. The power to identify the population structure can be evaluated from the quality of the plot and the accuracy of the discrimination. The power to retrain the geography or origin of the individuals can be assessed from the distance of the population and the match of populations to the corresponding regions in the scatter plot, as well as the correlations between the *F_ST_* and the Euclidean distance of the genetic features between each pair of population centroids.

### Application of ALFDA to the 1000 Genomes Project dataset

To test the ability of our approach to identify the population structure, we applied our method to 1000 genomes dataset containing 2504 individuals from 26 populations [51]. Datasets were downloaded from 1000 Genomes Project Phase III, which is available at https://www.internationalgenome.org/data. These datasets represent the worldwide populations from five continents. The population information including the abbreviations of the 26 populations and 5 super-populations as well as their collecting sites is summarized in Appendix 2 Sheet1.

To ensure the genotype matrix computed from the autosomes represent genome-wide structure. We carried out a Hardy-Weinberg equilibrium filter at *P* value < 0.001 and genotyping rate filter at > 0.02. We removed Major Histocompatibility Complex (MHC) regions on chromosome 6 and inversion region in the chromosome 8 as both were previously reported to have a large impact on population structure [52]. The remaining SNPs were pruned using PLINK 1.9 [53] with windows of 1000 variants and step size 10, pairwise squared correlation threshold at 0.02, and minor allele frequency > 0.05 to obtain SNPs in linkage equilibrium. Finally, we generated a genotype matrix with 2504 individuals carrying 38, 936 SNPs for population structure inference.

We carried out PCA analysis on the genotype matrix with genotype matrix scaled. The PCA scores were calculated and the first three PCA components were used to visualize the population structure of 26 populations. DAPC was then conducted using the first 20 PC scores labeled by the 26 populations. We conducted ALFDA on the genotype matrix with the *u* tuned from 0.1-10 (0.1, 0.5, 1, 5, 10). The first three reduced discriminant features were visualized as the representation of population structure.

Likewise, we performed *t-*SNE with its parameters tuned using the same procedure in simulation studies as described above. The optimal projection was chosen to visualize the population structure. UMAP was implemented with the number of nearest neighbours tuned from 20 to 99 (20, 40, 60, 80, 99) and the minimum distance tuned from 0.05 to 0.99 (0.05, 0.1, 0.5, 0.99). The best projection was chosen as the representation of population structure.

We plotted the first two reduced features as the representation of population structure for five approaches. We also attached the interactive plots for readers to look into the details of each projection in the appendix files.

### Application of ALFDA to 11, 640 Chinese individuals from the CONVERGE project

Despite rapid progress on genomic study in diverse populations, current existing genomic research still largely focused on populations of European descent. Populations from Asian, especially for the largest population in the world, Han Chinese, are still understudied. We applied our approach to visualize the population structure of the world largest ethnic group – Han Chinese populations. The dataset is obtained from the CONVERGE project (China Oxford and Virginia Commonwealth University Experimental Research on Genetic Epidemiology) [54] and the details of the dataset were described in [55]. The dataset consists of whole-genome sequencing (WGS) of 11, 640 women individuals (after QC filtering) collected from 58 hospitals in 45 cities and 24 out of 33 administrative divisions (19 provinces, 4 municipalities, and 1 autonomous region) across China. This dataset covers one of the broadest sampling of Chinese across China at present. We applied these five approaches to this data to compare their ability towards distinguishing large numbers of individuals at province level (including 4 municipalities and 1 autonomous region). The name of each province and their abbreviations are shown in Appendix 2.

We removed individuals that are ethnic minorities such as GuangxiZhuangzu. We removed the biallelic SNPs with the minor allele frequency (MAF) being smaller than 5%, and a missing rate of 5%. We did a Linkage Disequilibrium (LD) based SNP pruning with the threshold of *r*^2^ = 0.2. We kept 9,703 individuals belonging to 17 provinces for analysis.

PCA was carried out on genotype matrix after SNPs were filtered according to the above criteria. DAPC was then applied using the first 20 PCs produced from PCA which account for 9% genetic variance but no longer add discriminatory power to differentiate populations when increase more PCs. Likewise, t-SNE and the UMAP were implemented with their parameters tuning under a dense grid (the same procedure as above) and the best projections were chosen for population structure visualization. ALFDA was carried out accordingly with the optimal model was chosen after a dense parameter tuning on *u* with constant *K* equal to 3 (one less than the minimal number of individuals per population).

### Application to ancient individual genomes in Britain during the Middle to Late Bronze Age

We applied our approach to the ancient individuals from European countries (mainly Britain and Europe countries) during the Middle to Late Bronze Age (4000 BCE-43 CE) reported in Patterson et al. 2021. The raw data are available as aligned sequences (bam files) through the European Nucleotide Archive under accession number PRJEB47891. The datasets contain 826 individuals belonging to 19 countries sequenced by 1, 240K capture. The 1, 240K capture includes approximately 1.24M SNP sites in diverse modern and ancient populations [56]. The compiled datasets that passed QC, as well as the individual geographic coordinates and its country of collection are downloaded from the supplementary file of Patterson et al. 2021. In this study, we focus on 6 populations (Wales, Netherlands, Scotland, Channel Islands, England, France) consist of 511 samples around Britain. The geographic locations of the collected individuals are presented in Fig. S1 and Fig. 4A. We applied five methods to the genotypes of these samples and obtained the reduced features for visualizing the population genetic structure. PCA was carried out on the datasets with the minor allele frequency (MAF) being greater than 0.01, and the individual genotyping missing rate less than 50%. We did a Linkage Disequilibrium (LD) based SNP pruning with the threshold of *r*^2^ = 0.2. The first 50 PCs were used in DAPC analysis which account for 93% of the genetic variance. t-SNE and UMAP, and ALFDA were implemented with the same procedures as described in CONVERGE data analysis.

### Data availability

The simulated datasets in *gds* format are available in Zenodo at https://doi.org/10.5281/zenodo.7486563. The 1000 Genomes Project Phase III datasets are available at https://www.internationalgenome.org/data. The CONVERGE data are available at the European Nucleotide Archive (ENA) under accession number PRJNA289433. The raw data of the ancient genomes of Britain are available as aligned sequences (bam files) through the European Nucleotide Archive under accession number PRJEB47891.

### Code availability

Command line for simulations, scripts for simulation data analysis and empirical data analysis are available in Zenodo at https://doi.org/10.5281/zenodo.7486621. The ALFDA package is available at https://github.com/xinghuq/ALFDA.

### Ethics declarations

### Ethics approval and consent to participate

Ethical approval was not needed for this study.

## Acknowledgments

XQ is supported by International Postdoctoral Exchange Fellowship Program (Introduction Program, YJ20210310) and the General Program of China Postdoctoral Science Foundation (2022M723075) as well as the CAS Special Research Associate Funding. PJ is funded by the Strategic Priority Research Program of the Chinese Academy of Sciences (XDB38010400), National Natural Science Foundation of China (32270706). The authors thank Prof Oscar Gaggiotti for valuable comments.

## Author Contributions

XQ designed the study, carried out the analyses and wrote the article with the supervision of PJ. All authors read, edited, and approved the submitted version of the manuscript.

