## Supplementary Materials for "New machine learning method identifies subtle fine-scale genetic stratification in diverse populations"

**Supplementary Information**

**Supplementary Tables**

Table S1. Simulation parameters of four demographic models

| **Scenarios** | **Regions** | **Number of populations** | **Population size (diploid)** | **Sample size** | **Migration rate** | **Recombination rate/gen/bp** | **Mutation rate** | **Number of markers/per chromosome** |
| --- | --- | --- | --- | --- | --- | --- | --- | --- |
| **Island model** | 4 (4,4,4,4) | 16 | 1000 | 100 | 0.001 | 1×10^-8^ | 1×10^-8^ | 100,000 |
| **Hierarchical island model** | 4 (4,4,4,4) | 16 | 1000 | 100 | *m*_within_: 0.001 | 1×10^-8^ | 1×10^-8^ | 100,000 |
|  |  |  |  |  | *m*_between_: 0.0001 |  |  |  |
| **Stepping stone** | 2 (8,8) | 16 | 1000 | 100 | 0.001 | 1×10^-8^ | 1×10^-8^ | 100,000 |
| **Hierarchical stepping stone** | 2 (8,8) | 16 | 1000 | 100 | *m*_within_ : 0.001 | 1×10^-8^ | 1×10^-8^ | 100,000 |
|  |  |  |  |  | *m*_between_: 0.0001 |  |  |  |

**Supplementary Figures**

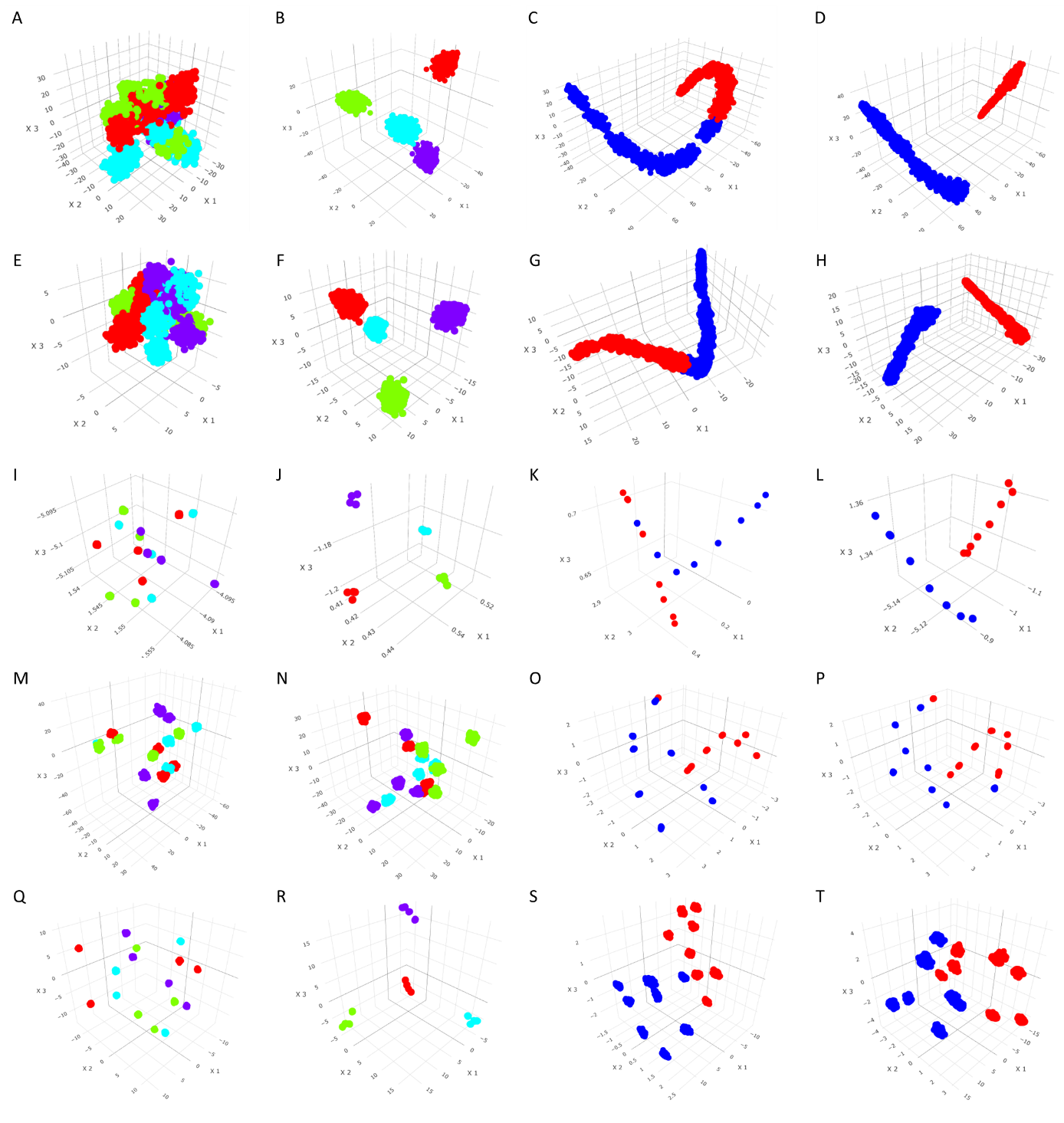

Fig. S1. 3D visualization of the reduced genetic features under four spatial scenarios by different approaches (A, E, I, M, Q: island model; B, F, J, N, R: hierarchical island model; C, G, K, O, S: stepping stone model; D, H, L, P, T: hierarchical stepping-stone model) using PCA, DAPC, ALFDA, t-SNE and UMAP. A-D, Genetic structures of four spatial scenarios inferred from PCA; E-H, Genetic structures of four spatial scenarios inferred from DAPC; I-L, Genetic structures of four spatial scenarios inferred from ALFDA. M-P, Genetic structures of four spatial scenarios inferred from t-SNE. Q-T, Genetic structures of four spatial scenarios inferred from UMAP. The same colour in the scatter plots represents the same region.

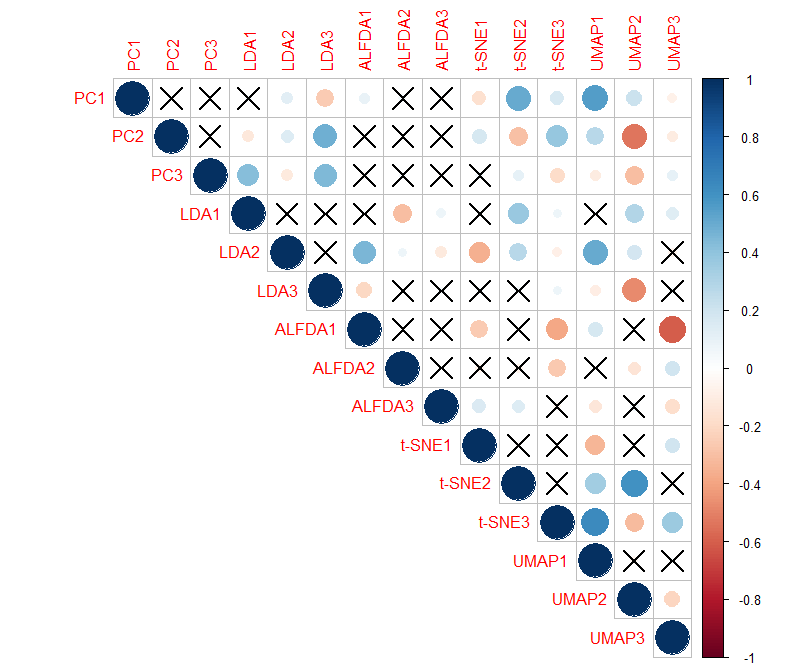

A

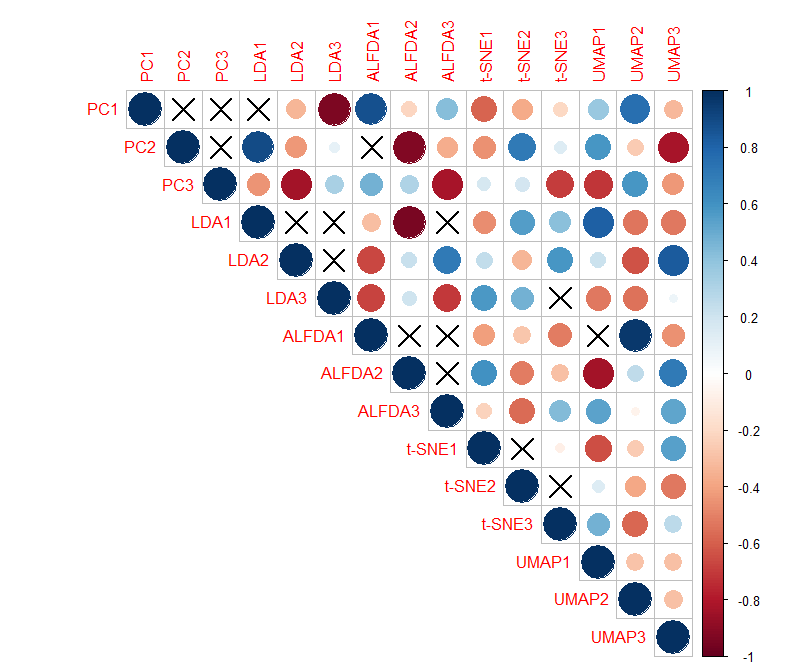

B

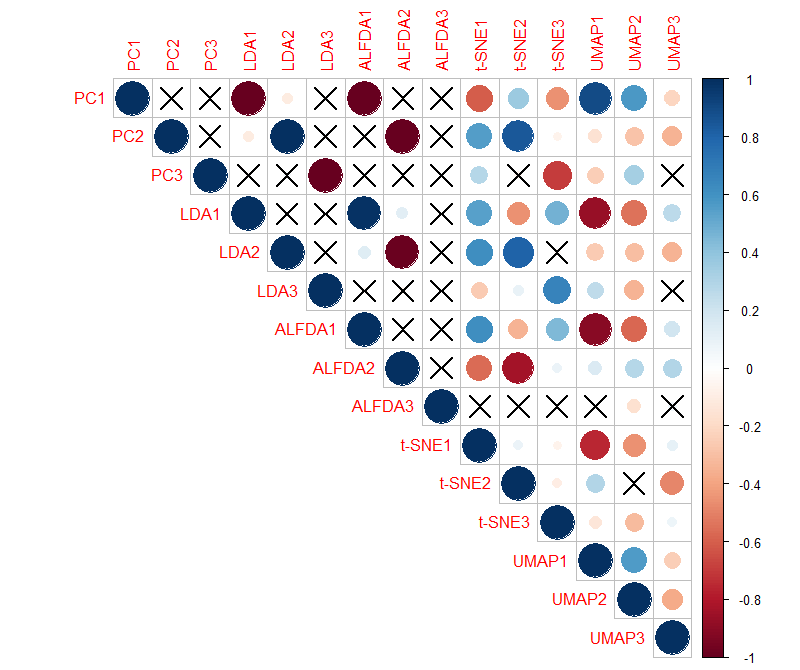

C

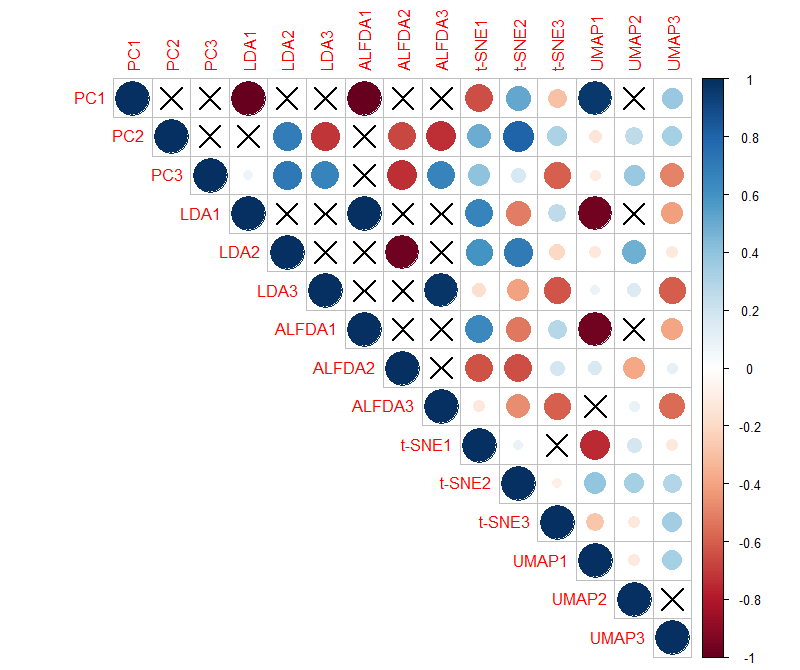

D

Fig. S2. Correlations between the first three reduced features from five approaches under (A) Island model, (B) Hierarchical Island model, (C) Stepping stone model, and (D) Hierarchical stepping stone model. Non-significant correlations (P value larger than 0.5) are marked as ⨯.

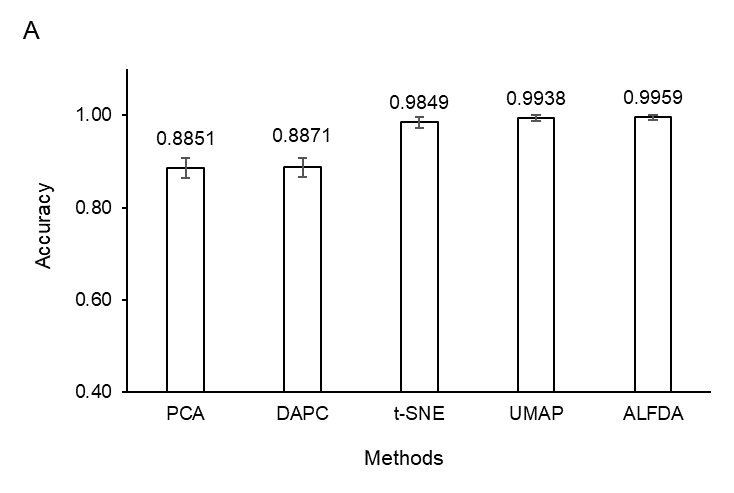

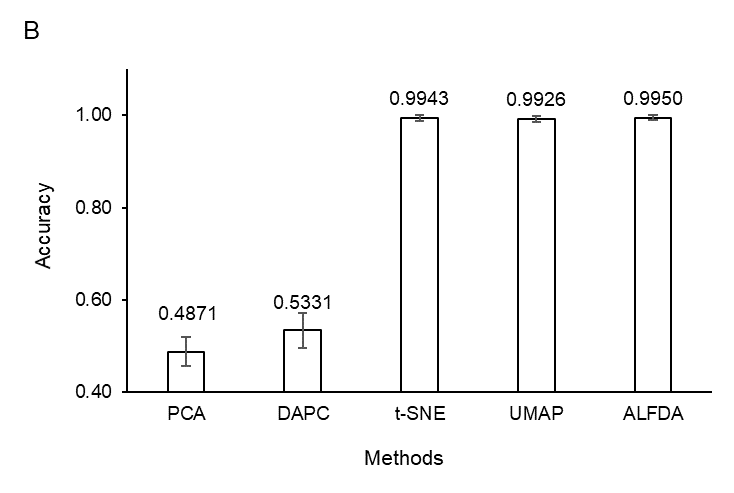

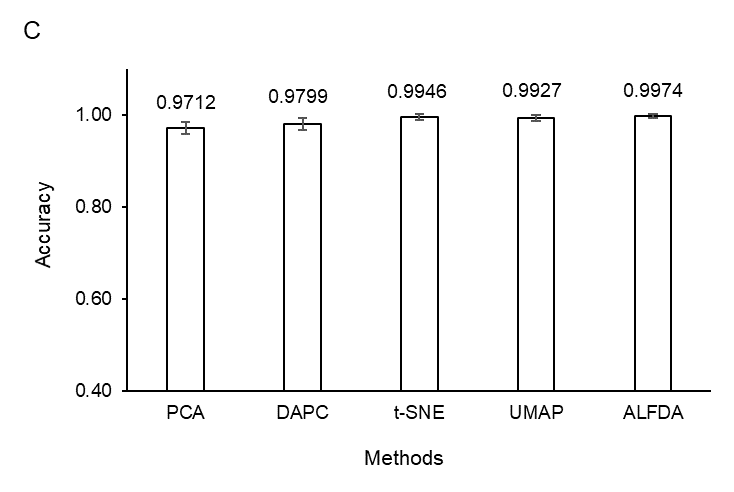

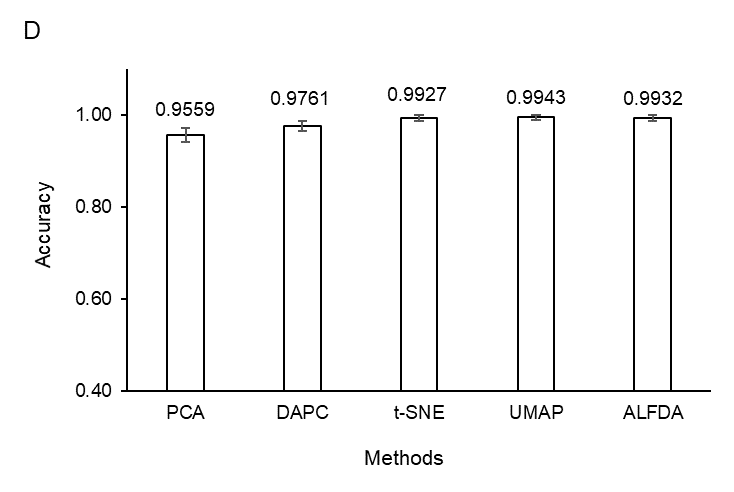

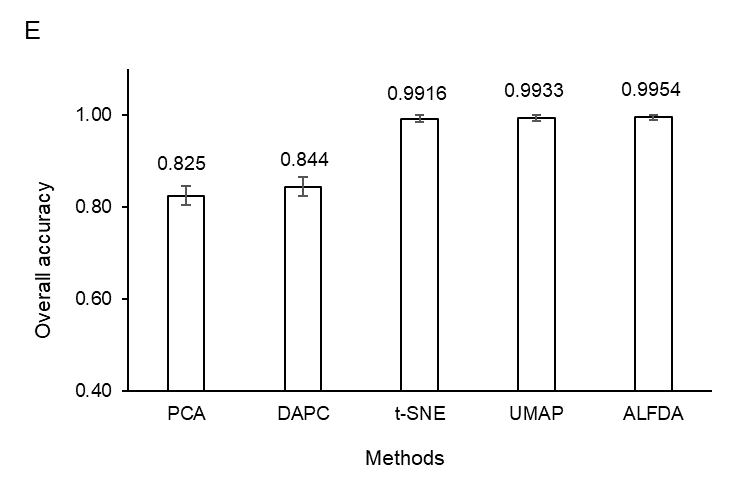

Fig. S3. Discriminatory power of five machine learning approaches in distinguishing populations under (A) Island model, (B) Hierarchical Island model, (C) Stepping stone model, (D) Hierarchical stepping stone model, as well as their overall accuracy (E).

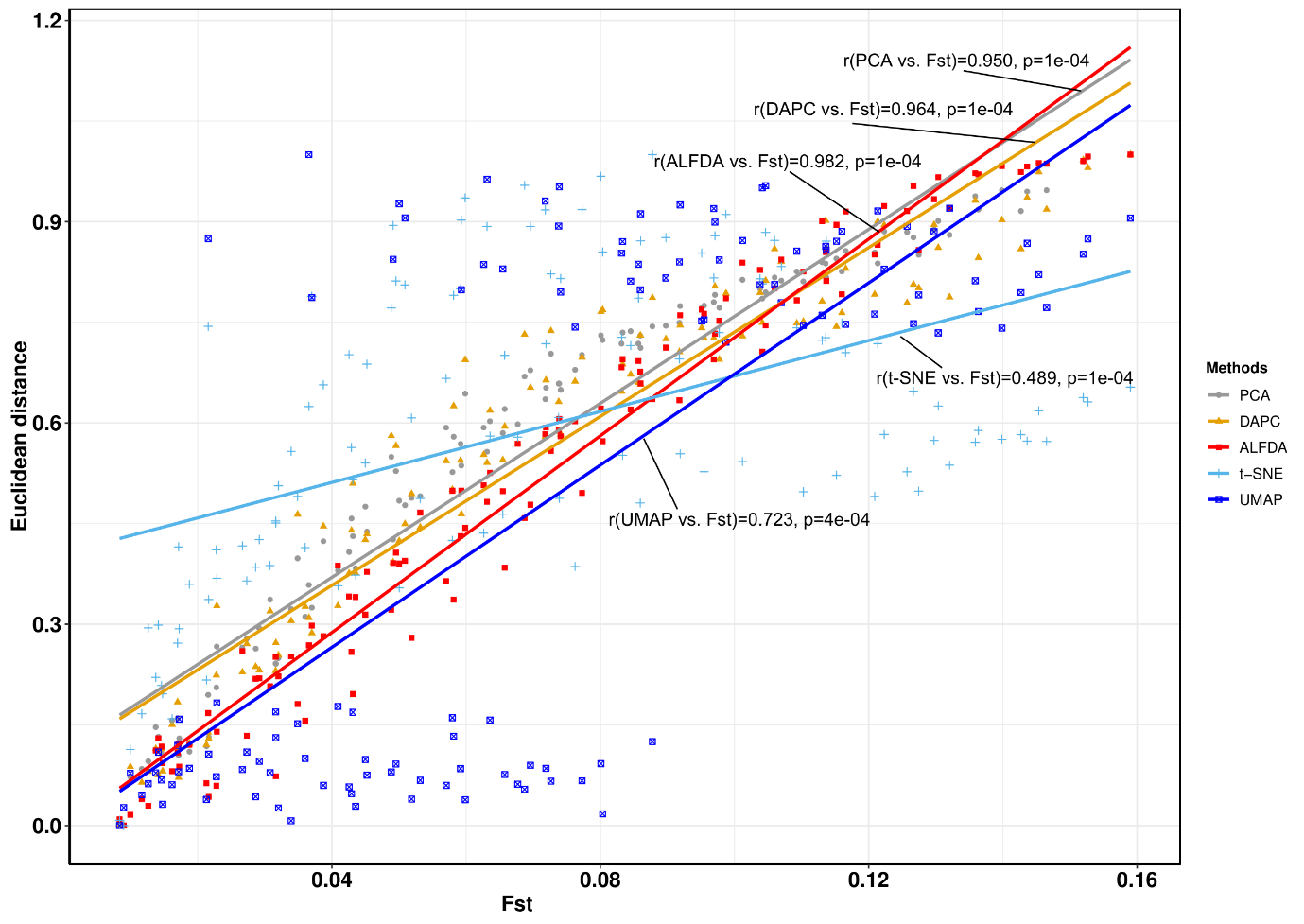

Fig. S4. The Mantel's r between the population genetic fixation Fst and the Euclidean distance of the centroid of clusters projected by five methods under isolation by distance model. All distances are scaled to 0–1.

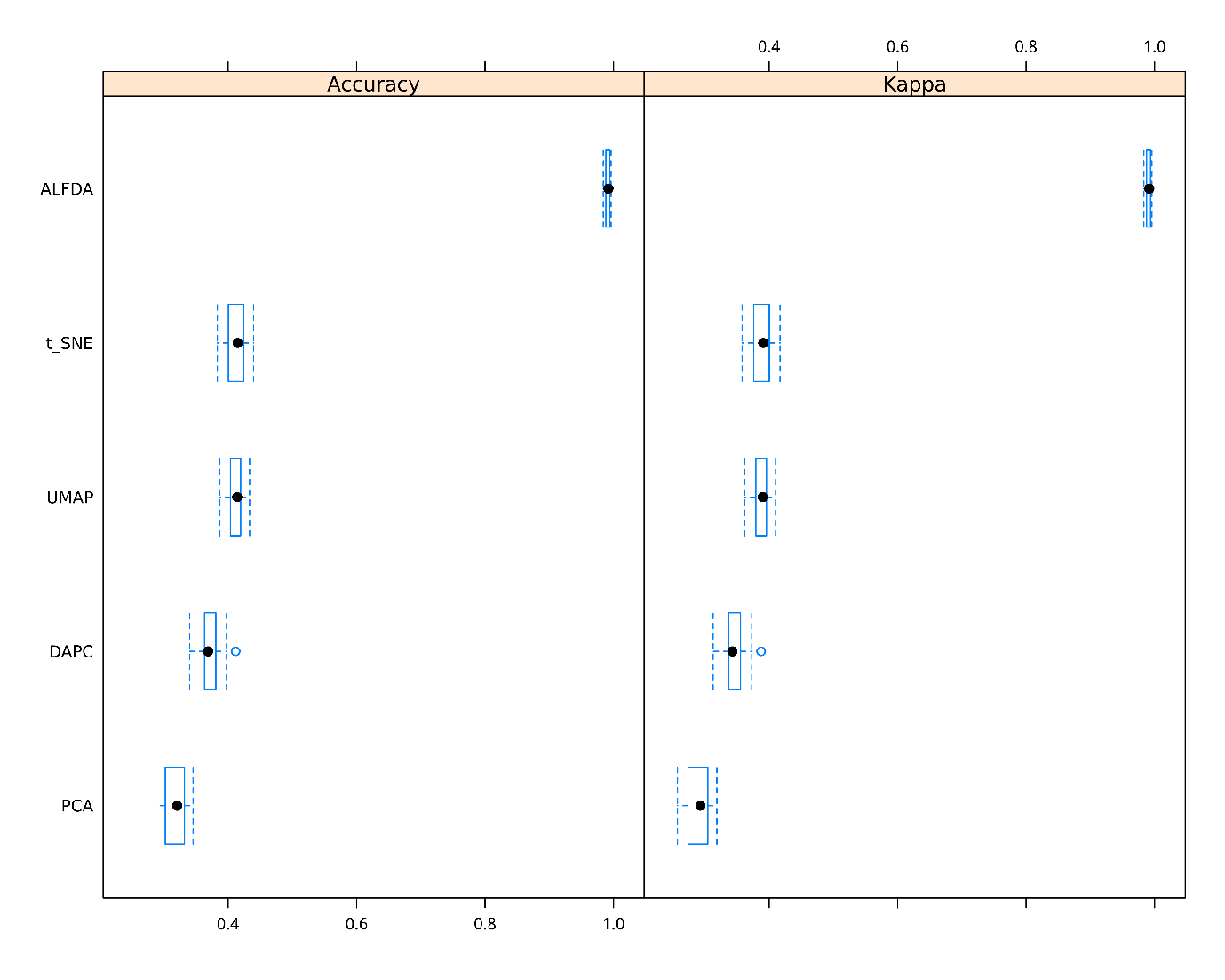

b

d

d

c

c

b

b

b

a

a

Fig. S5. The performance (accuracy and Kappa values) of five machine learning approaches to assign individuals to their corresponding countries in 1000 genomes dataset. The accuracy and Kappa values were calculated using the average values of 100 resamples (training and testing) obtained from cross validations.

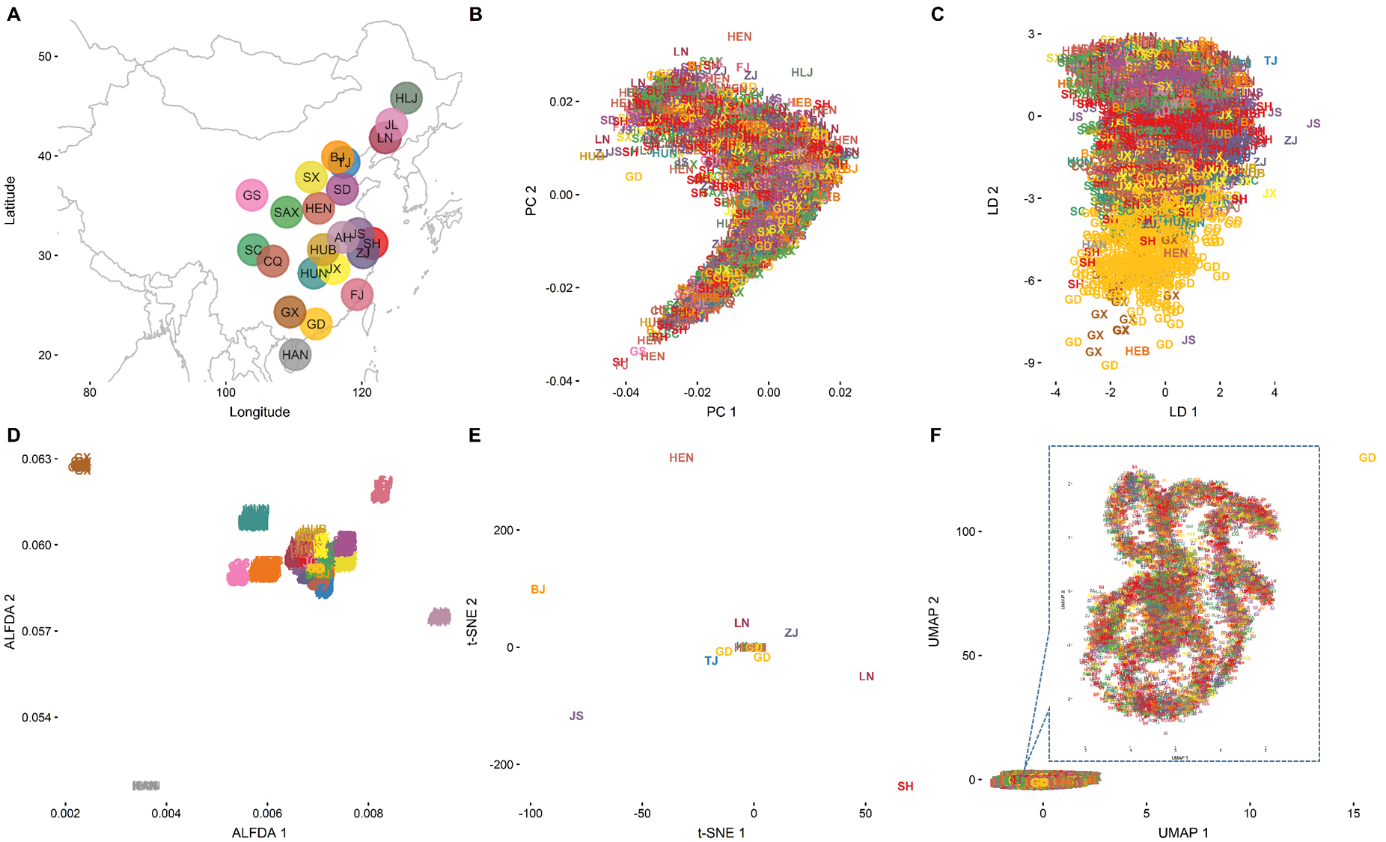

Fig. S6. (A). The geographical locations of each province of China. Population structure of Chinese women from 24 provinces represented by (B) PCA, (C) DAPC, (D) ALFDA, (E) t-SNE, and (F) UMAP. Individual labels were assigned based on their self-reported birth locations and colored according to the map in figure (A). The PCA, DAPC and ALFDA representations were projected after a 90° clockwise rotation. Province abbreviations: Shanghai, SH; Liaoning, LN; Zhejiang, ZJ; Tianjin, TJ; Hunan, HUN; Sichuan, SC; Shaanxi, SAX; Heilongjiang, HLJ; Jiangsu, JS; Shandong, SD; Henan, HEN; Hebei, HEB; Beijing, BJ; Guangdong, GD; Jiangxi, JX; Shanxi, SX; Hubei, HUB; GuangxiZhuangzu, GX; Chongqing, CQ; Fujian, FJ; Gansu, GS; Jilin, JL; Anhui, AH; Hainan, HAN.

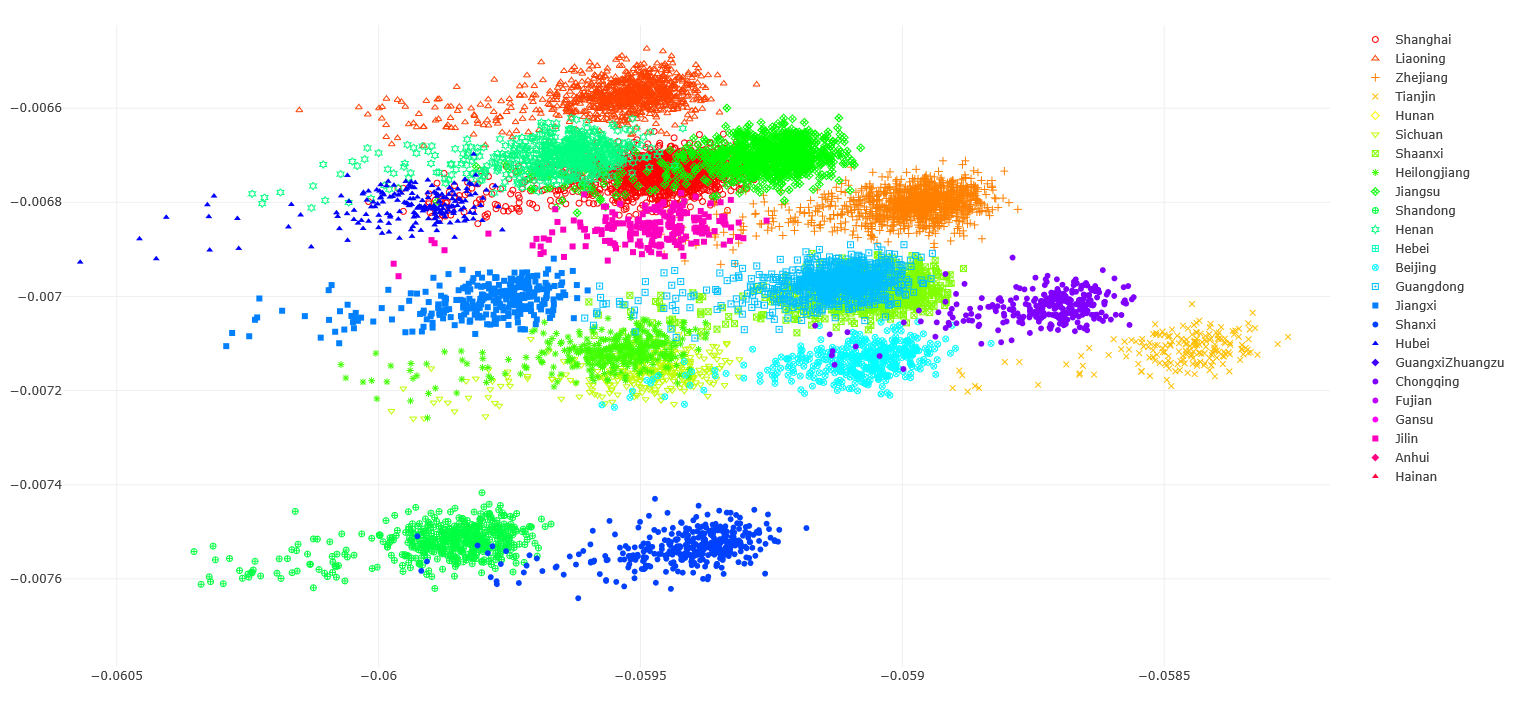

Shanghai

Fig. S7. Population structure at the province level with the specific focuses on the closed populations from Fig. S7. Populations from Anhui, Hainan, Hunan, Gansu, and Hebei are excluded.

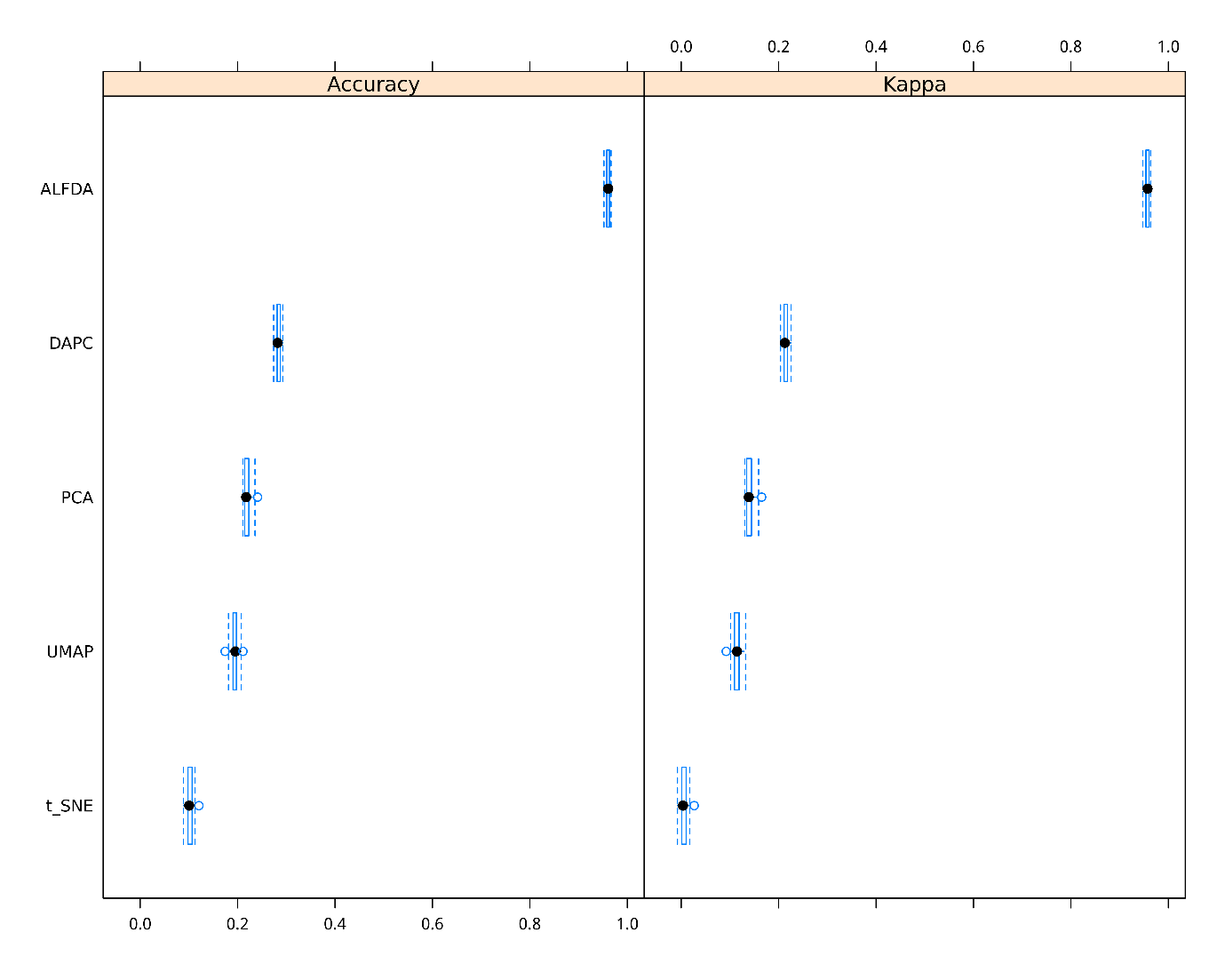

e

e

d

d

c

c

b

b

a

a

Fig. S8. The performance (accuracy and Kappa values) of five machine learning approaches to assign individuals to their self-reported birth provinces in CONVERGE dataset. The accuracy and Kappa values were calculated using the average values of 100 resamples (training and testing) obtained from cross validations.

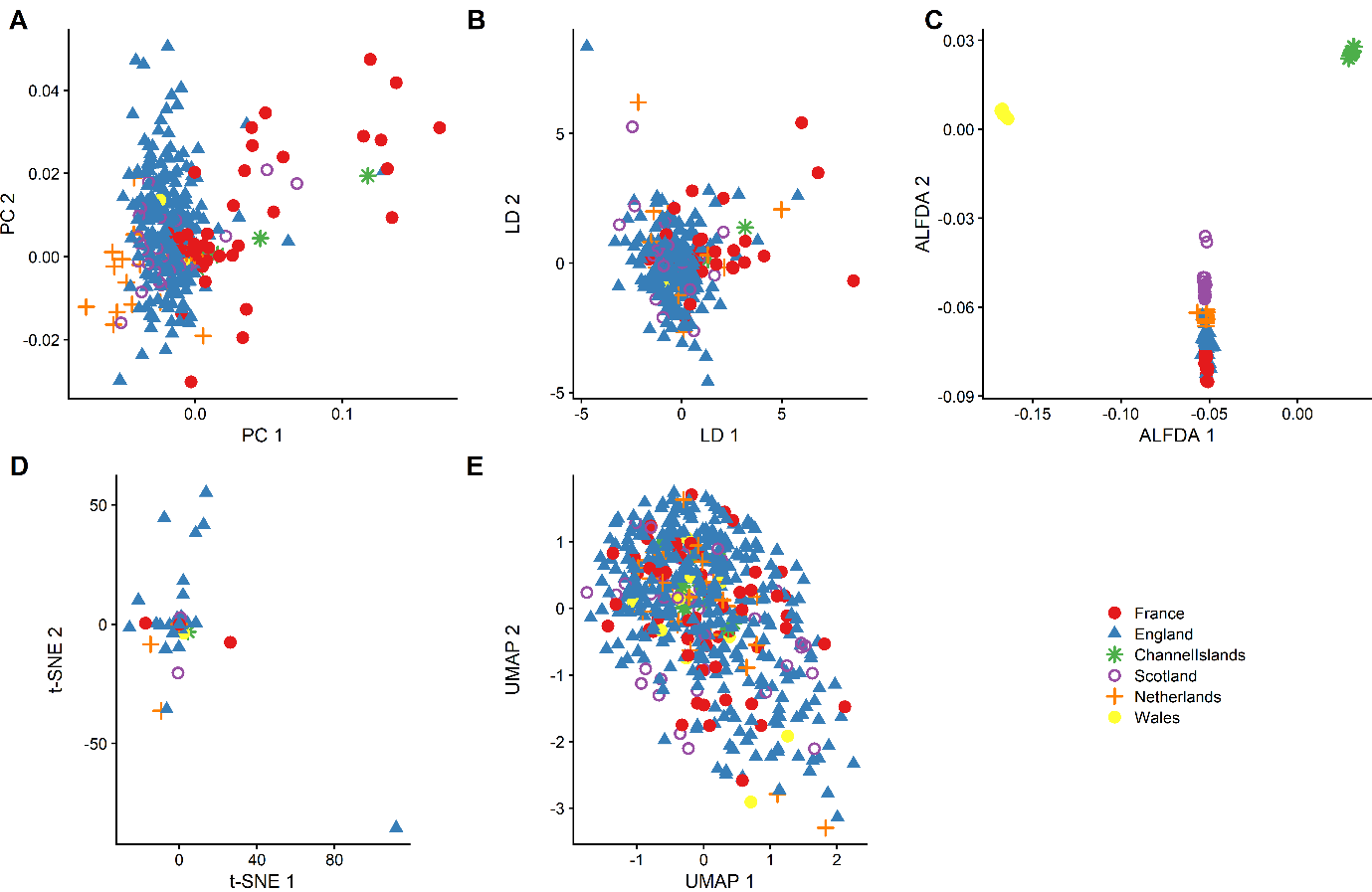

Fig. S9. Population structure of four populations from Britain and Europe. Population genetic structure projected by PCA (A), DAPC (B), ALFDA (C), t-SNE (D), UMAP (E).

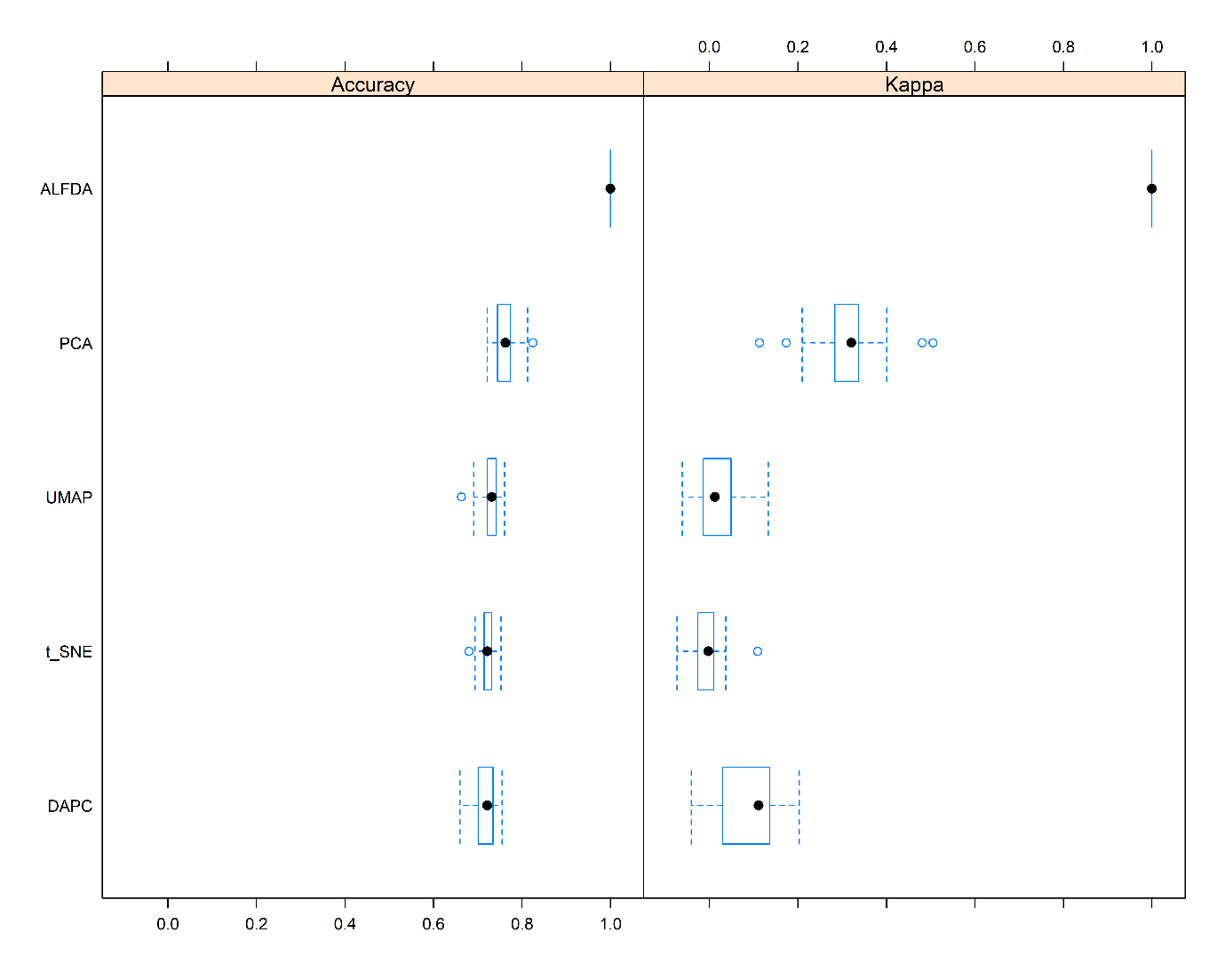

a

b

c

c

c

c

c

b

b

a

Fig. S10. The performance (accuracy and Kappa values) of five machine learning approaches to assign individuals to their sampled countries based on the dataset obtained from Patterson et al., 2021. The accuracy and Kappa values were calculated using the average values of 100 resamples (training and testing) obtained from cross validations.

**Supplementary Methods**

***Locality-Preserving Projection***

Locality-Preserving Projection seeks to find the local structure using the nearest neighbor algorithm. Assuming$\bar{A}_{i,j}^{s}$ is the affinity matrix between two individuals (samples) *x_i_* and *x_j_*. $\bar{A}_{i,j}^{s}$ belongs to (0,1], the closer the two points, the larger the $\bar{A}_{i,j}^{s}$. Let *x_i_* ϵ *N^d^* (*i* = 1, 2,…, *n*) be *d*-dimensional samples and *y_i_* ϵ (1,2, …, *g*) be associated class labels, where *n* is the number of samples and *g* is the number of classes. Let *z_i_* ϵ *N*^m^ (1 ≤ *m* ≤*d*) be embedded samples, where *m* is the dimension of the embedding space. *Z_i_* can be obtained using the *d ⨯ m* transformation matrix *z_i_*=*T^T^ x_i_*.

The LPP transformation matrix can then be defined as the following:

$T_{LPP}={argmin}_{T\epsilon R^{d⨯m}}\frac{1}{2} \sum_{i,j=1}^{n} \bar{A}_{i,j}^{s}{|\left| T^{T}x_{i}-T^{T}x_{j} \right||}^{2}$ ,	(S1)

subject to $T^{T}$*X*$\bar{D}$*X* $T^{T}$=$\bar{I}$,	(S2)

where *X* is the matrix of all samples, *X* = (*x_1_|x_2_|…|x_n_*); $\bar{I}$ is the identity matrix and $\bar{D}$ is the diagonal matrix with *i*-th diagonal element being

 $\bar{D}_{i,i}=\sum_{i=1}^{n} \bar{A}_{i,j}^{s}$ .	(S3)

As shown in equation (S1) and (S2), *T* is the transformation matrix that keeps the nearby data pairs close in the embedding space.

Let ${[\alpha_{i}]}_{i=1}^{d}$ be the generalized eigenvectors associated with the generalized eigenvalues λ1 ≥ λ2 ≥ … λ*d* of the following generalized eigenvalue problem:

*X*$\bar{L}$*X*^T^*α* = *λX*$\bar{D}$*X*^T^*α*,	(S4)

where $\bar{L}$*=*$\bar{D}$*-*$\bar{A}^{s}$ .	(S5)

The solution of the LPP transformation (S3) is given by

*T_LPP_* = (α*_d_*|α*_d_*_-1_| … |α*_d_*_-_*_r_*_+1_).	(S6)

***Non-linear local Fisher discriminant analysis***

To measure differences between populations while also capture the local variances, we use the ratio of between-population and within-population variances, also known as the *F-* statistic to find the best direction *T* to maximize Fisher’s criterion:

*F*_(_*_T_*_)_ =$\frac{{T^{T}S}^{(b)}T}{{T^{T}S}^{(w)}T}$ .	(S7)

*F*_(_*_T_*_)_ is the ratio of between-population affinity and within-population affinity, which can be used to define the extent of differentiation between populations, with larger values indicating stronger differentiation between populations. $S^{(b)}$ and $S^{(w)}$ represent the between population scatter matrix and within population scatter matrix respectively. Below, we present the non-linear local Fisher discriminant analysis model in using to extract the population structures.

Let $\bar{x}_{i}$, and $\bar{x}_{j}$represent the *i-*th and *j*-th samples. $\bar{A}_{i,j}^{s(w)}$ represents the within- population affinity and $\bar{A}_{i,j}^{s(b)}$ represents the between-population affinity. Let $\bar{S}^{m}$ be the local mixture scatter matrix defined by $\bar{S}^{m}$ = $\bar{S}^{(w)}$ + $\bar{S}^{(b)}$. The local within-population scatter matrix $\bar{S}^{(w)}$ and the local between-population scatter matrix $\bar{S}^{(b)}$ can be obtained as follows.

$\bar{S}^{\left( m \right)}$= $\frac{1}{2}$ $\sum_{i,j=1}^{n} \bar{A}_{i,j}^{s\left( m \right)} {\left( x_{i}-x_{j} \right)\left( x_{i}-x_{j} \right)}^{T}$ ,	(S8)

$\bar{A}_{i,j}^{s\left( m \right)}$=$\bar{A}_{i,j}^{s\left( w \right)}+\bar{A}_{i,j}^{s\left( b \right)} = \left\{ \begin{aligned} \frac{{\bar{A}^{s}}_{i,j}}{n} , \mathrm{if}y_{i} = y_{j} \\ \frac{1}{n} , \mathrm{if}y_{i} \neq y_{j} . \end{aligned} \right\}$ .	(S9)

Therefore, $\bar{S}^{\left( m \right)}$ is equal to

$\bar{S}^{\left( m \right)}$=$\frac{1}{2}$ $\sum_{i,j=1}^{n} \bar{A}_{i,j}^{s\left( m \right)} (x_{i}{x_{i}}^{T}+x_{j}{x_{j}}^{T}-x_{i}{x_{j}}^{T}-x_{j}{x_{i}}^{T})$

=$\sum_{i=1}^{n} (\sum_{j=1}^{n} \bar{A}_{i,j}^{s\left( m \right)} )x_{i}{x_{i}}^{T}-\sum_{i,j=1}^{n} \bar{A}_{i,j}^{s\left( m \right)}x_{i}{x_{j}}^{T}$.	(S10)

This can be expressed in matrix form as

$\bar{S}^{m}=X\bar{L}^{(m)}X^{T}$ , (S11)

where $\bar{L}^{(m)}$ =${\bar{D}^{\left( m \right)}- \bar{A}}^{s\left( m \right)}$ and $\bar{D}^{\left( m \right)}$ is the *n*-dimensional matrix with the *i-*th diagonal element being $\bar{D}_{i,i}^{\left( m \right)}$=$\sum_{j}^{n} \bar{A}_{i,j}^{s\left( m \right)}$.

Likewise, $\bar{S}^{(b)}$ can be obtained by $\bar{S}^{(b)}$= $X\bar{L}^{(b)}X^{T}$, where $\bar{L}^{(b)}$=$\bar{D}^{\left( b \right)}-\bar{A}^{s\left( b \right)}$, and $\bar{D}^{\left( b \right)}$ is the *n*-dimensional matrix with the *i-*th diagonal element being $\bar{D}_{i,i}^{\left( b \right)}$=$\sum_{j}^{n} \bar{A}_{i,j}^{s\left( b \right)}$.

The computation of the matrix, $\bar{S}^{(w)}$ and $\bar{S}^{(b)}$, can be achieved by eigenvector decomposition, $\bar{S}^{(m)}V$=$\lambda\bar{S}^{w}V$, which can be expressed as

$X\bar{L}^{(m)}X^{T}V$=$\lambda X\bar{L}^{(w)}X^{T}V$,	(S12)

For any vector *V* belonging to $R^{d}$ can be expressed by using some vector *α* belonging to $R^{n}$ as *V* = *X^T^α*. Therefore, $X^{T}V$ =${XX}^{T}V\alpha$=*K*α. Then, multiplying Eq. (S12) by $X^{T}$ from the left-hand side, we have

${M\bar{L}}^{(m)}M\alpha$= $\lambda M\bar{L}^{(w)}M\alpha$ ,	(S13)

where *M* is an *n*-dimensional matrix (*n* ⨯ *n*) with the (*i*, *j*)-th elements being $M_{i,j}$=$x_{i}^{T}x_{j}$. Until now, we have successfully non-linearized the local Fisher discrimination. The local linear discriminant problem converts to be the distance $M\left( x_{i},x_{j} \right)$ discrimination problem.

Let $M\left( x_{i},x_{j} \right)$ be the distance between sample $x_{i}$and $x_{j}$. $M_{i,j}$ ϵ *N^n^*^⨯^*^n^* (*i* = 1, 2, …, *n*) is *n*-dimensional distance matrix and with sample label *y_i_* ϵ (*C*_1_, *C*_2_, …, *C_g_*). Let $x_{g}^{i}$ be the sample *x_i_* belonging to population *C_g_*, with population size being *N_g_*. Therefore, the within-population scatter matrix $\bar{M}^{(w)}$ and between-population scatter matrix $\bar{M}^{(b)}$ can be defined as

*m_av_*= ($\frac{1}{n}$ $\sum_{i=1}^{n} M\left( x_{1},x_{i} \right),$ $\frac{1}{n}$ $\sum_{i=1}^{n} M\left( x_{2},x_{i} \right),\ldots, \frac{1}{n}$ $\sum_{i=1}^{n} M\left( x_{n},x_{i} \right)$ )*^T^*,	(S14)

*m_i_*= ($\frac{1}{N_{i}}$ $\sum_{g=1}^{N_{i}} M\left( x_{1},x_{g}^{i} \right),$ $\frac{1}{N_{i}}$ $\sum_{g=1}^{N_{i}} M\left( x_{2},x_{g}^{i} \right),\ldots, \frac{1}{N_{i}}$ $\sum_{g=1}^{N_{i}} M\left( x_{n},x_{g}^{i} \right)$),	(S15)

$\bar{M}^{(b)}$ =$\sum_{i=1}^{g} \frac{N_{i}}{n} \left( m_{i}-m_{av} \right)\left( m_{i}-m_{av} \right)^{T}$,	(S16)

$\bar{M}^{(w)}$=$\frac{1}{n}\sum_{i=1}^{g} \sum_{g=1}^{N_{i}} (M_{i,+}-m_{i}){(M_{i,+}-m_{i})}^{T}$,	(S17)

Although the kernel version of local Fisher discriminant analysis (KLFDA) employing kernel matrix has successfully improved the discrimination accuracy, choice of the optimal kernel function and their parameters are still tricky, and there is no simple generalization to multi-class discrimination. Furthermore, KLFDA suffers from data piling because of the large diagonal values (diagonal dominance) when dealing with large Gram matrix [1]. We solve these problems and improve the non-linear data mapping by introducing of the weighted local affinity matrix. The distance matrix *M* is the weighted genetic affinity matrix, $M_{i,j}$= ${A_{g}^{us}}_{i,j}$. Then, the final *F* statistic now becomes

$F_{(T)}$ =$\frac{{T^{T}M}^{(b)}T}{{T^{T}M}^{(w)}T}$ .	(S18)

The finally eigenvalue solution becomes $\bar{M}^{(b)}$𝛼 = 𝜆$\bar{M}^{(w)}$𝛼. In practice, the matrix $\bar{M}^{(b)}$ is often singular. A regularized method is often used to solve the problem, which is transformed into a general eigenvalue problem by choosing a smaller positive number *θ*, in which case can be expressed as

${(\bar{M}}^{(w)})^{-1}\bar{M}^{(b)}$𝛼 = 𝜆𝛼,

$\bar{M}^{(w)}=\bar{M}^{(w)}+\theta\bar{I}$,

$(\bar{M}^{(w)}+\theta\bar{I})^{-1}\bar{M}^{(b)}$𝛼 = 𝜆 ,	(S19)

where *θ* is the regularization parameter, $\bar{I}$ is the identity matrix. The performance of the discrimination is dependent on *θ*. The recommended default *θ* value is 0.001. However, the optimal *θ* value can be also determined through the empirically estimation [2] ,

$\bar{M}^{(w)}$+*θ*$\bar{I}$ →0 .	(S20)
